# Reduction in RBD Binding Affinity to Glycosylated ACE2 is Entropic in Origin

**DOI:** 10.1101/2022.10.12.511994

**Authors:** Mauro L. Mugnai, Sucheol Shin, D. Thirumalai

## Abstract

The spike protein in the virus SARS-CoV-2 (the causative agent of COVID-19) recognizes the host cell by binding to the peptidase domain (PD) of the extracellular enzyme Angiotensin-converting Enzyme 2 (ACE2). A variety of carbohydrates could be attached to the six asparagines in the PD, resulting in a heterogeneous population of ACE2 glycoforms. Experiments have shown that the binding affinity of glycosylated and deglycosylated ACE2 to the virus is virtually identical. In most cases, the reduction in glycan size correlates with stronger binding, which suggests that volume exclusion, and hence entropic forces, determine the binding affinity. Here, we quantitatively test the entropy-based hypothesis by developing a lattice model for the complex between ACE2 and the SARS-CoV-2 spike protein Receptor-binding Domain (RBD). Glycans are treated as branched polymers with only volume exclusion, which we justify using all atom molecular dynamics simulations in explicit water. We show that the experimentally measured changes in the ACE2-RBD dissociation constants for a variety of engineered ACE2 glycoforms are well accounted for by our theory, thus affirming that ACE2 glycans have only a weak, entropic effect on RBD binding.

## Introduction

Glycosylation, a common post-translational modification, is characterized by the attachment of oligosaccharides (glycans) to specific locations on a protein.^1^ We focus here on N-linked glycosylation, in which short carbohydrates are covalently bound to the side-chain of asparagines at the beginning of a N-X-S/T sequon, with X being any residue except proline. ^1^ Glycoproteins occur in a variety of glycoforms: an amenable asparagine may or may not be glycosylated, and the oligosaccharide found at a given site may feature different sugars, various degrees of branching, and chains of different lengths. ^2^ Both computational and experimental studies have illustrated that glycosylation stabilizes the glycoprotein against unfolding^3,4^ and aggregation.^2,5^ In particular, larger glycans reduce protein aggregation propensity more than smaller ones, which hints at a prominent role played by volume exclusion and, hence entropic forces.^5^ However, proteins may also specifically bind glycans.^6^ Such interactions often play a role during the initial step of viral invasion, involving the recognition of the host cell.^7^ The receptors used by viruses to attach to the host include carbohydrates forming the glycocalyx surrounding the cell.^7^ Here, we focus on the role of some glycans in the initial recognition of human cell by the virus SARS-CoV-2, which is the causative agent of COVID-19.

The spike protein of the virus SARS-CoV-2 is a homotrimer decorated with a variety of glycans.^8–10^ Remarkable molecular dynamics (MD) simulations have shown that the SARS-CoV-2 glycans are flexible, and cover a large fraction of the spike surface.^11,12^ Their presence is thought to hide the epitopes from the immune system.^8,12,13^ The name “glycan shield” (or equivalently glycan camouflage) vividly illustrates this concept.

In the first step of viral invasion, the SARS-CoV-2 spike protein exposes the receptor-binding domain (RBD),^14^ which engages with an extracellular protein anchored to the cell membrane called Angiotensin-converting Enzyme 2 (ACE2).^15,16^ The peptidase domain (PD) of ACE2 (residues 1-614) includes the RBD binding site, and the 6 asparagines that are susceptible to glycosylation: N53, N90, N103, N322, N432, and N546.^9,17,18^

An important question to answer is whether the glycans associated with ACE2 strengthen or weaken RBD binding to the human receptor. In other words, does the spike protein specifically recognize the glycans on ACE2? Atomically detailed MD simulations ^9,19–26^ have found energetic interactions between ACE2 oligosaccharides N-linked to N53, N90, N103, N322, or N546 with the RBD and its N-linked glycan at N343. In particular, some of these studies suggest that the N90 glycan protects ACE2 from RBD binding.^20,21,23^ Accordingly, a deep mutational scanning of ACE2 revealed that mutations that disrupt the N90-X-S/T sequon increase RBD binding affinity.^27^ In contrast, the N322 glycan was suggested to strengthen the interaction between the virus and ACE2.^20,21^ In addition, it was predicted that different glycans linked to a given site may either stabilize or destabilize the complex.^23^ Force-pulling simulations showed that ACE2 glycans extend the range of interaction between ACE2 and the RBD,^26^ and reduce the complex dissociation rate measured by Atomic Force Microscopy.^24^ From the MD studies, the contacts between ACE2 glycans and the RBD were identified,^20–22,24,26^ and the associated energy of glycan-RBD interactions was calculated.^20,21^ In addition, by studying the conformations of the ACE2 glycan in the absence of RBD, MD simulations report on the likelihood that a oligosaccharide “shields” the RBD-binding site on ACE2.^20^ Finally, MD simulations can identify whether the presence of the RBD modifies the extent of glycan fluctuations.^25^ Although most interesting, these findings provide only a qualitative picture. In particular, the relation between observables reported in the MD studies and experimentally measurable quantities, such as the free energy of binding, was not clear. Nevertheless, the paradigm emerging from at least some of these studies^20,21,23,24^ seems to be that ACE2 glycans are strong regulators of ACE2-RBD binding, and that the details of glycosylation site and the structure of the oligosaccharide chain modulate the interactions in unique ways.

In apparent contrast to the conclusion from MD simulations, experiments have shown that favorable energetic interactions involving glycan with ACE2 do not regulate binding to RBD. *In vitro*^25^ and *in vivo*^28^ studies showed that partial or even full deglycosylation of ACE2 has only a small impact on protein stability. Thus, glycan engineering could be used to reveal the importance of carbohydrates in ACE2-RBD binding. Measurements of the ACE2-RBD dissociation constant revealed that modifying the glycoform of ACE2 has minimal impact on RBD binding.^18,25,26,28–31^ Upon complete (or nearly complete) removal of one or more glycans, in almost all cases (with only one exception, referring to a dimeric ACE2 construct^31^) the dissociation constant changes by less than a factor of 2 to 3 as compared to the wild type (WT). Such a small change corresponds to a standard free energy change of ⪅ 1*k*_B_*T*. In most cases, it was found that partial or complete deglycosylation increased binding affinity or had a very small effect^18,25,29,30^ (note that some authors report that ablating the ACE2 glycans results in increased dissociation constant compared to the WT^26,32^). In addition, as illustrated with the moss plot representation,^12^ solvent exposed glycans appear to be generally flexible,^11,12,20,21^ although the exact extent of their fluctuations depends on their local environment.^11^ These observations prompted us to hypothesize that the effect of ACE2 glycans on RBD binding should be predominantly entropic in origin with the energetic interactions playing a sub-dominant role. The physical picture is that RBD binding sterically reduces the number of conformations accessible to the glycans (Fig. 1). This is in agreement with the role that glycans play in reducing the propensity for proteins to aggregate.^5^ In the context of RBD-ACE2 binding, others have put forward similar ideas following computational,^20,25^ theoretical, ^33^ or experimental^29^ studies. Here, using lattice models supported by MD simulations, we show that glycan entropy suffices to account for the impact of ACE2 glycans on RBD-ACE2 binding. Our calculations explain the current experimental data nearly quantitatively.

**Figure 1:**
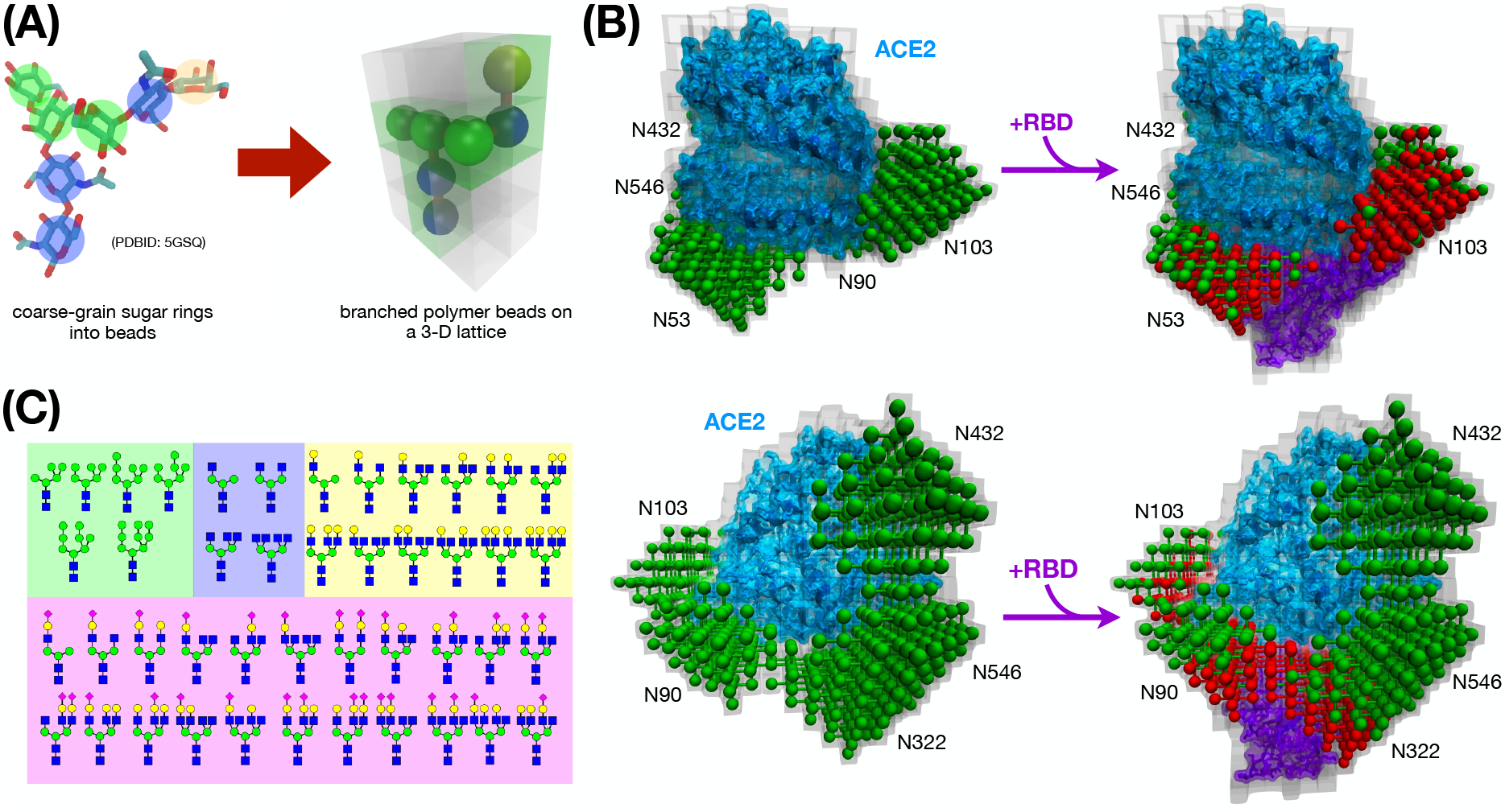
Construction of the model. (A) Schematic illustration of a coarse-grained glycan, modeled as a branded polymer whose beads represent the individual sugar rings (left). The lefthand beads-on-string polymer is put on a three-dimensional cubic lattice to fluctuate and occupy the volume in space with its given configuration (right). (B) Graphical representation for how the glycans in ACE2 are affected by RBD binding in our model. Glycans attached to N53, N90, N103, N322, N432, and N546 are shown as beads and sticks within the cubic lattice sites (gray shades). On the lefthand side, the oligosaccharides shown in green represent the lattice-space ensemble of their configurations that do not overlap with the lattice sites occupied by ACE2 (cyan; from PDBID: 6LZG). On the righthand side, the glycan configurations that are found to clash with the lattice sites occupied by RBD (purple; also from PDBID: 6LZG) are portrayed in red. The bottom panel is the same as the top, except that it is rotated by about 180 degrees. (C) All the glycans considered in this study. Different sugars are colored and shaped according to convention (blue squares for N-acetylglucosamine, green circles for mannose, galactose is shown as yellow circles, and sialic acid is reported as a magenta diamond). The background color corresponds to the color of the last saccharide in the chain, and it is used in Fig. 4 to distinguish the various oligosaccharides. In the model, all sugars are indistinguishable and interact solely by volume exclusion.

## Results

### Glycan Model

We developed a three-dimensional cubic lattice model of the glycoprotein ACE2 and its interaction with the RBD. In this representation, glycans are modeled as branched polymers on a lattice, where each lattice site roughly represents the center of a sugar ring (see Fig. 1A). We should note that the description of glycans as polymers on a lattice was introduced some time ago to investigate how oligosaccharides modulate glycoprotein folding.^34^ Using the lattice representation of glycans, we quantify how the ensemble of the glycan conformations changes upon the RBD-ACE2 binding (Fig. 1B; more details to be described below). Figure 1C shows a list of the various glycans considered in this study, which vary in size (degree of polymerization, or number of monosaccharides composing the chain) and connectivity (branching). For each glycan in Fig. 1C, we sample *exhaustively* the ensemble of all self-avoiding conformations (see Supporting Information, section).

In order to account for the geometry of glycans, the lattice spacing is chosen to be *a* = 5.5 Å, which is approximately commensurate with the distance between the centers of mass of two consecutive sugar rings along the oligosaccharide chain obtained using atomistic MD simulations (Fig. 2A). The details of the MD simulations are provided in section of the Supporting Information (SI). We tested whether the self-avoiding lattice model provides a reasonable description of the size of the glycans, presumed to be flexible in our model, by comparing the distribution of the end-to-end distance (Fig. 2B) and the radius of gyration (Fig. S6) with the MD simulation result. The distributions obtained for the lattice model are broader and with smaller averages, indicating that the discretized glycans are more compact than the more realistic off-lattice, all-atom counterparts. It is likely that the recovery could be improved by introducing sugar-sugar repulsion beyond volume exclusion; however, this would detract from the simplicity of the model. The difference between the average of the distributions could be reduced by using a larger value of the lattice spacing. As we discuss later, we carried out such a test and we found out that the results did not change the essential conclusions of our work. In light of these considerations, we deem the discrepancies between the distributions to be acceptable.

**Figure 2:**
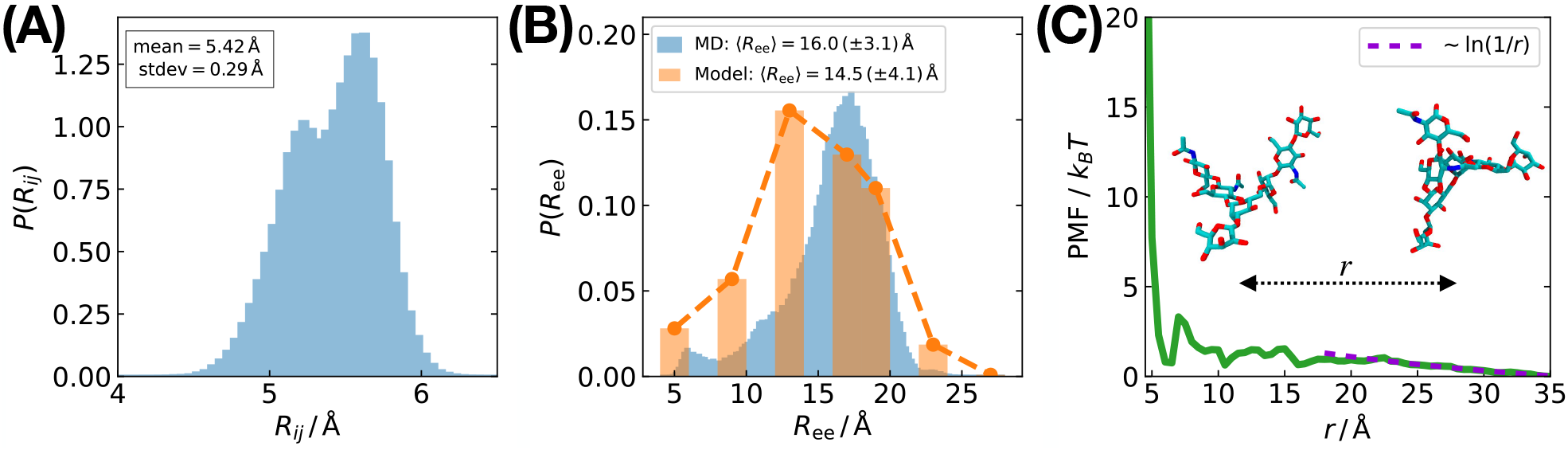
All-atom MD simulations reveal that glycans are flexible and soluble in water. (A) Distribution of the distance between the center of mass of consecutive sugar rings obtained using the MD simulation. (B) Comparison between the distributions of the end-to-end distance computed from the MD simulation and our lattice model. (C) Potential of mean force (PMF) for a pair of the glycans as a function of the pair distance, *r*, where the dashed line is the fitting curve for large *r*.

### MD simulations for Inferring Glycan Interactions

In our model, we render the interactions within a glycan, between glycans, or involving glycan and protein to be repulsive (volume exclusion), which is tantamount to assuming that glycans have no energetic preference for being fully exposed to water or for interacting with the protein surface or other glycans. This approximation was initially suggested by the portrayal of the glycan shield conformations through MD simulations, ^11,12^ which indicated that within the MD timescale solvent exposed glycans did not appear to be adsorbed on the protein surface. This is in line with the intuitive expectation that sugars dissolve well in water. Nevertheless, to test and quantify this assumption, we explored the interactions between glycans using explicit-solvent, atomistic MD simulations. By performing umbrella sampling calculations^35^ for a pair of identical glycans, we determined the potential of mean force (PMF) between them (see section of the SI for details). As shown in Fig. 2(C), overall the PMF increases upon decreasing the pair distance, *r*. When the glycans are close enough to be in contact (*r* < 15 ≈ 2*R*_*g*_, where *R*_*g*_ ≈ 7.5 is the radius of gyration of the glycan) the PMF increases rapidly (Fig. 2(C)) due to volume exclusion. We note parenthetically that the PMF decays as ∼ ln(1*/r*) for large *r*, which is reminiscent of the effective interaction between two polymer-grafted colloidal particles in a good solvent.^36^ Although there is a shallow basin at *r* = 6.5, the depth is only about 2 *k*_*B*_*T*, which is not large enough to maintain a stable dimer. Direct simulations of glycan aggregation support this observation. We simulated four identical glycans in a periodic box of various sizes, which changes the concentration of the carbohydrates. We found that the glycans can form a transient dimer but the pair contact probability and aggregation probability among more than three glycans are proportional to [glycan] and [glycan]^2^ (where [glycan] = *N*_glycan_*/L*^3^, with *N*_glycan_ being number of oligosaccharides in a cubic periodic box of side *L*), respectively, which are the scaling relations expected for an ideal solution (see Fig. S8). Therefore, the small attractive interaction between glycans does not lead to the formation of stable aggregates. We confirmed that the simulated systems are equilibrated, by calculating the ergodic measure, ^37,38^ see Fig. S8. Thus, we conclude that overall a lattice model and volume exclusion provide a simple but reasonable description of the glycans.

### Glycan Impact on ACE2-RBD Binding

First, we mapped the structure of the ACE2-RBD complex to the lattice (Fig. 1B). In order to do so, we first centered the structure around asparagine *ν* ∈ 𝒩, where 𝒩 = {N53, N90, N103, N322, N432, N546}. Second, we selected all of the sites on the lattice that are within *a/*2 + *r*_ha_ from an ACE2 or RBD heavy atom (C, N, O, or S) of van der Waals radius *r*_ha_.^39^ The sites that satisfy this distance criterion are occupied by the protein complex. Third, for each glycan *g*, we counted the number 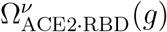 of oligosaccharide conformations that do not clash with the protein complex. Finally, we removed the RBD from the ACE2-RBD complex, and we repeated the calculation, which yields the number of self-avoiding conformations 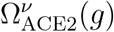. Under the approximations that (i) the proteins do not move, (ii) glycans bound to different sites are independent, and (iii) the effect of glycans in the RBD can be ignored, we calculated the dissociation constant using,

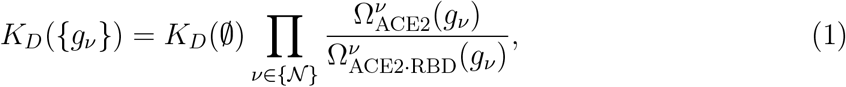

where {*g*_*ν*_} refers to a set of glycans (one glycan per glycosylation site), and *K*_*D*_(∅) is the dissociation constant in the absence of glycans. In order to account for the heterogeneity of the glycoforms reported for ACE2, the measured dissociation constant becomes,

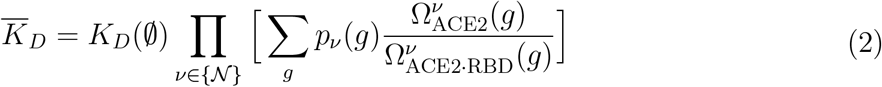

where *p*_*ν*_(*g*) is the probability that glycan *g* is N-linked to asparagine *ν* (Σ_*g*_ *p*_*ν*_(*g*) = 1 for each asparagine), with the sum covering the whole ensemble of possible oligosaccharides. In practice the summation is limited to those portrayed in Fig. 1C, including the possibility that the site is empty (*g* = ∅). Equation (2) follows from Eq. (1) after minor manipulations with the assumption that the glycans occupying different sites are independent. Note that in Eqs. (1)-(2) the contribution of the glycans to the dissociation constant is exclusively entropic in origin. Although partially justified using MD simulations, the assumptions of the theory, leading to Eq. (2), can only be assessed by comparing the predictions with experiments. (For further details on the algorithm, and for the derivation of Eqs. (1)-(2) see the Supporting Information sections -. Note that, in the Supporting Information, we use the over bar to denote an average over the glycoforms, which we omit for the sake of simplicity in the main text.)

### Summary of Experimental Results

Before assessing the validity of our theory, we summarize some key experimental findings. A number of published experimental results report on the dissociation constants between the RBD (or the spike protein) and either the WT ACE2 or specific glycoforms obtained by adopting protocols that alter the glycan population.^18,25,26,30,31^ For instance, it is possible to use (i) enzymes that fully (or almost completely) remove the N-linked oligosaccharides,,^18,25,26,30^ (ii) or that increase the presence of sialic acids (+ST6^18^). In addition, (iii) sialidases (+sia) are adopted in order to remove sialic acids,^18,25^ and (iv) kifunesine ensures that only mannose-type carbohydrates are linked to the protein (+Kif). Moreover, ACE2 can be altered to remove specific glycans, for instance the T92Q^25^ and T92I^31^ mutations preclude glycosylation at asparagine N90, and the N322Q mutant^25^ prohibits the attachment of an oligosaccharide to asparagine N322. Finally, we ignore results obtained upon removal of fucoses, because ablation of these sugars has little effect on the RBD dissociation constant,^18,32^ likely due to their distance from the proteinprotein binding interface. ^32^

### Comparison with Experiments

We compared our theoretical predictions with the dissociation constants measured for all of the above ACE2 glycoforms after scaling the experimental value by either the dissociation constant for WT ACE2 [*K*_*D*_(WT)], or by *K*_*D*_(∅), which was given by the dissociation constant obtained after partially or fully ablating the glycans. We computed the model dissociation constant using Eq. 2 with *p*_*ν*_(*g*) adapted from those provided either by Crispin and coworkers^18^ or by Azadi and coworkers^17^ (this required small manipulation of tabulated or plotted data; for further details, see Supporting Information section). Figure 3 compares the results of our calculations and experiments. Clearly, the dissociation constants are in the right range with the agreement being very good over-all. The trends from experimental data are captured, although the model shows a smaller decrease in *K*_*D*_ after ablation of sialic acids or for constructs lacking the N90 glycosylation site.

**Figure 3:**
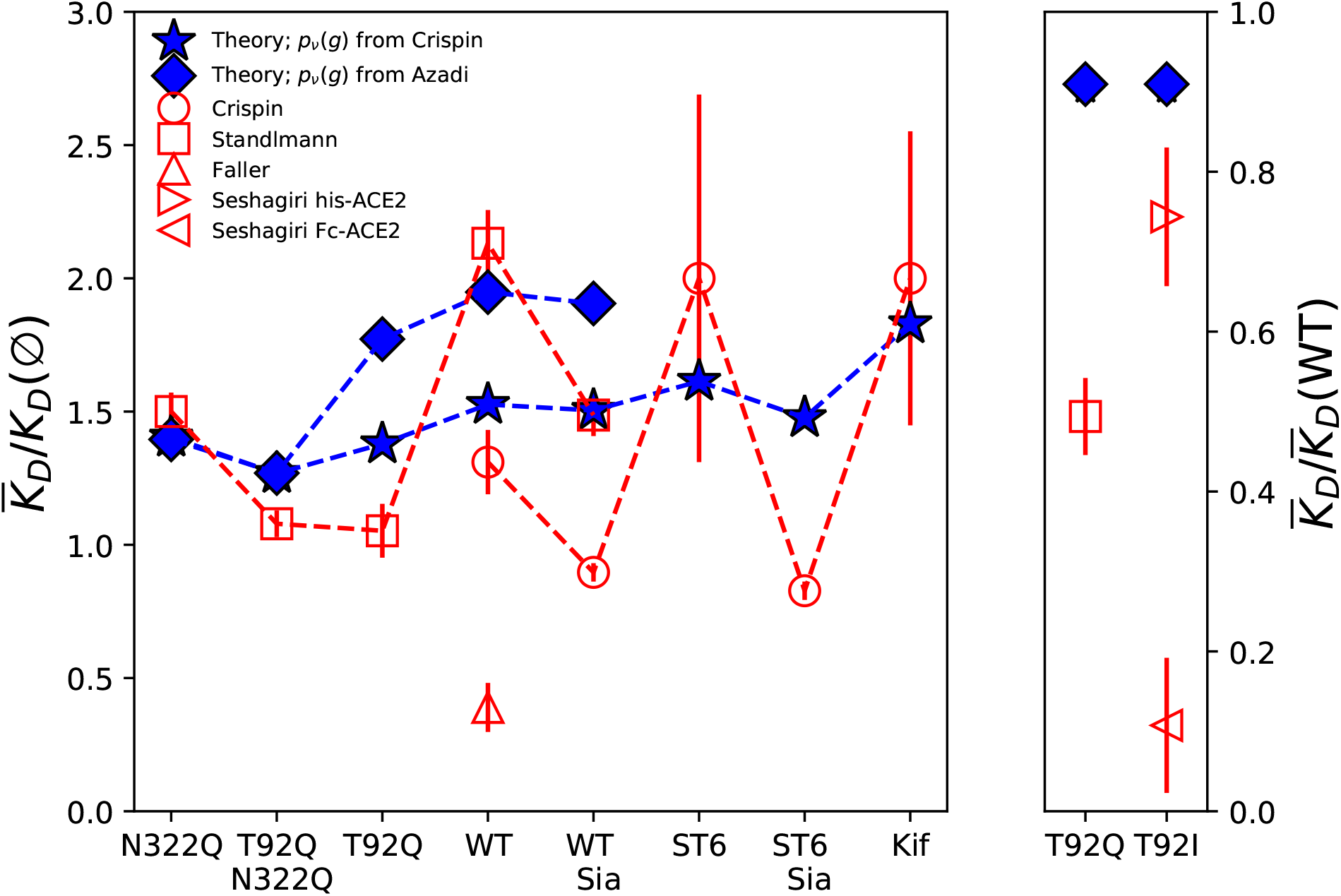
Comparison between experiment and the theoretical predictions. The x-axis refers to different modifications of the ACE2 glycoforms through sequence mutation (T92Q, N322Q, T92I) or enzymes (ST6, Sia, and Kif). The y-axis on the left panel is the ratio between a dissociation constant and the value obtained from the same experiment when all of the glycans are ablated (in some cases only a sugar is left). In the right panel, the y-axis shows the ratio between the dissociation constant between modified and WT ACE2. The red symbols are from experiment: circles for Crispin and coworkers,^18^ squares for Standlmann and coworkers,^25^ up-pointing triangles for Faller and coworkers,^26^ and right- and left-pointing triangles for the experiments conducted by Seshagiri and coworkers^31^ with His and Fc tagged ACE2, respectively. The blue stars (diamonds) show the results from the model, and they are obtained using the glycan distribution from Crispin and coworkers^18^ (Azadi and coworkers^17^).

Next, we calculated the contribution to the dissociation constant of each oligosaccharide in our ensemble at the six ACE2 sites considered (see Fig. 1B). For each of the four sites, Ω_ACE2_*/*Ω_ACE2·RBD_ depends linearly on the degree of polymerization (Fig. 4). The results depend on the geometry of the complex, and the distance of the glycosylated sites from the binding interface. Clearly, the contribution of oligosaccharides bound to N103, N432, and N546 are negligible, as might be expected given that all of these oligosaccharides are N-linked to ACE2 that are far away from the RBD. Three asparagines, N53, N90, and N322, can occupy the volume taken by the RBD in the complex, and affect the binding to RBD according to the following series: N322 *>* N53 *>* N90. It is unclear if this order is supported by experiments. Deep mutagenesis^27^ has provided evidence that the removal of the N90 glycan increases the affinity for ACE2. Indeed, experiments report that the N90X (with X any residue), I91P, and S92X (with X any residue other than T) are all favorable for RBD binding. The results for N53 and N322 are not clear because the only mutations reported that impact glycosylation at these sites are I54 and T344. I54P should eliminate the N53 glycan, and has a small effect on binding. T344S increases binding without affecting glycosylation, indicating that other effects on the structure of ACE2 might also contribute.

**Figure 4:**
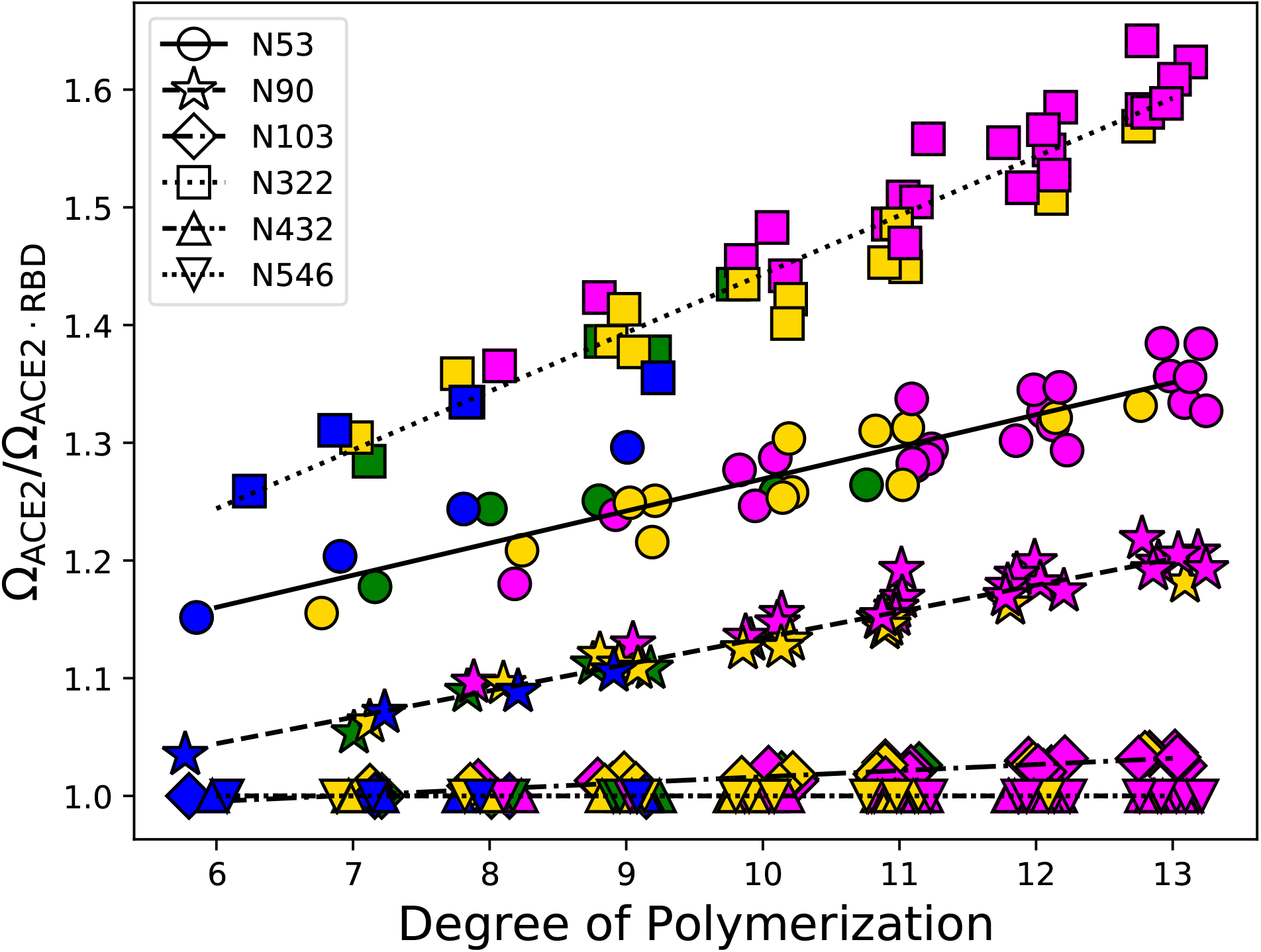
Contribution of all glycans to the dissociation constant for RBD-ACE2 binding from theory. The y-axis shows the ratio between the number of self-avoiding glycan conformations N-linked ACE2 in isolated ACE2 and in complex with RBD as a function of the degree of polymerization of the glycans. Different symbols indicate the asparagine to which a glycan is linked (see legend). Colors green, blue, yellow, and magenta are used to identify the last sugar of the chain, as shown by the background of Figs. 1D. Lines are fit to the data. Note that the data were subject to a small amount of random noise along the x-axis to aid visualization. The degree of polymerization is an integer in all cases.

We should point out that MD simulations have identified N90^20^ and N322^20,21^ as interacting strongly energetically with RBD in order to strengthen the complex, which is not supported by experiments unless there is a compensating entropy loss. Although our analysis focuses on entropic contribution alone, we can make the following argument to extract the binding energy. Assuming that our lattice model provides the *exact* entropic contribution to the binding free energy, then we could estimate the binding energy by taking the difference of the logarithm of *K*_*D*_*/K*_*D*_(∅) obtained in experiments (which of course includes energy) and our model (which accounts only for entropy). Because the experimental results and our model are within a factor 2 from each other (with a sign that depends on each case), we can estimate that the energy of relative energy of binding |ΔΔ*H*| ⪅ *k*_*B*_*T* ln 2 ≈ 0.7*k*_*B*_*T*, a small contribution, in apparent disagreement with some results obtained in MD simulations. Of course, it is worth pointing out that this argument is approximate, as it is tantamount to associating the same binding energy to all the glycan conformations (an approximation common to some mean-field models such as Bragg-Williams and Flory-Huggins^40^).

## Discussion

Despite the success in reproducing nearly quantitatively the outcome of a variety of experiments from different groups, the validity of our model may be legitimately questioned on the grounds that it is too simple. The major assumptions that we have made are the following:

1. we modeled glycans on a lattice and assumed that they interact only via volume exclusion;
2. we used a discrete lattice model to describe protein structure and glycan conformations;
3. the conformational fluctuations of the proteins are neglected;
4. glycans are treated as independent from each other. We conducted some tests to support these approximations and to explore their limitations. Of course, the final crucifer are experimental data, which is well accorded for by our theory. Our study leads to new insights and predictions.

1. The use of lattice models has provided testable theories in protein folding ^41^ and amyloid fibril growth,^42^ and has also been exploited to study glycoprotein folding.^34^ Nevertheless, the discretized representation of the conformational ensemble of the glycan is certainly approximate. Ultimately, our model relies on the assumption that although the number of conformations available to each glycan depends on the discretization on the lattice, the ratio (which is related to the dissociation constant, see Eq. 1) does not. In addition, although MD simulations substantiate our conjecture that glycans can be treated as self-avoiding polymers (see Fig. 2), it is likely that oligosaccharides could preferentially stick to certain areas of the protein compared to others. Therefore, we cannot exclude that a more sophisticated inter-play between enthalpy (absent from our model) and entropy (which is our model accounts for) would play a role in shaping the affinity between ACE2 and RBD. All-atom models are of course better suited to compute free energies. However, such simulations needed to compute the dissociation constant as we did here (including glycoform heterogeneity) would be extremely challenging due to a large enthalpy-entropy compensation that is required to reproduce the measured disassociation constants. Current efforts have mostly focused on energy,^20,21^ and although some studies did explore conformational flexibility,^25^ they have not reached the quantitative recapitulation of the experimental results shown here.
2. The discretized representation of the protein is also another approximation. The resolution of the model is given by the diameter of each sugar, *a* = 5.5 Å, which was extracted from the distance between consecutive sugar rings in a model glycan subject to MD simulations (Fig. 2A). It is worth emphasizing that *a* is the only parameter in the model, and its value was not chosen to optimize agreement with the experimental results. We tested the robustness of the model upon changes in the lattice spacing by repeating the calculations using different values of *a*. Figures S4A-B show that changing *a*, within a range that covers most of the distribution in Fig. 1C, has a negligible effect on the dissociation constant. Figures S4C-D illustrate that the relative importance of the various glycans depends on the lattice spacing, in particular the impact of the N53 glycan on the dissociation constant diminishes as *a* increases, whereas the opposite happens for the carbohydrates attached to N90 and N322. Nevertheless, oligomers attached to N53 and N322 have consistently a larger impact on the affinity between ACE2 and RBD than N90-bound glycans.
3. Protein fluctuations are neglected in this study. Protein flexibility is likely to alter the number of allowed glycan conformations. Consider glycan N90, whose contribution to the dissociation constant appears larger in experiments than predicted by the theory (Fig. 3). As an illustration, let us consider *a* = 5.0 Å. In the crystal structure, PDBID: 6LZG, the ACE2 lysine K26 forms a wedge that restrains the orientation of the first sugar ring of the N90 oligosaccharide (see Fig. S5A). This restraint is immutable in our model, but protein and side chain flexibility might soften this constraint and enhance the fluctuations of glycan N90. To test this assertion, we constructed the K26A ACE2 mutant by removing all the lysine side chains after the beta carbon (this is of course only an approximation: the K26A might have other, less trivial effects on the structure of the protein). Compared to the WT ACE2, the mapping of K26A on the cubic lattice resulted in only one fewer bead, adjacent to the location of the anchor for the N90 glycan (see Fig. S5B). This small difference increased the number of conformations for the N90 glycan, resulting in a stronger impact on the *K*_*D*_*/K*_*D*_(∅) ratio, as shown by the comparison of blue/yellow/magenta starts (WT) with the cyan stars (K26A) in Fig. S4C. We should note that with lattice spacing *a* = 5.5 Å the discretized WT and K26A ACE2 are identical, so this analysis does not hold at lower resolution, as one might expect. Overall, these observations highlight a limitation of the model.

### On the correlations between glycans and a testable prediction

(4) Treating the different N-linked glycans as independent is necessary if the focus is on exhaustive sampling, and on accounting for the heterogeneous family of oligosaccharides linked at the various ACE2 N-glycosylation sites. However, this neglects the possibility that nearby glycans could occupy each other’s space, thereby reducing the number of available conformations (“crowding effect”), which could impact the affinity of ACE2 for the RBD. The direct test of this effect requires exhaustive enumeration of the conformations explored by pairs of glycans, which is practically impossible and ultimately is the reason for assuming that glycans are independent. We found that the ratio between the number of conformations in which there is an overlap between glycans *g*_1_ and *g*_2_ N-linked to ACE2 glycosylation sites *ν*_1_ and *ν*_2_ 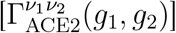 and the number of conformations of the two glycans assumed to be independent 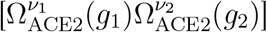 is bound by,

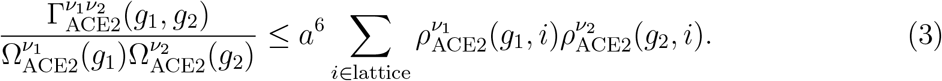

In Eq. (3), *i* is the index that identifies a vertex in the cubic lattice, and 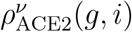 is the density at site *i* of the glycan *g* attached to site *ν*. The r.h.s. is a measure of the overlap between the two densities. The formal proof of Eq. (3) is in the Supporting Information section. The argument leading to the bound in Eq. (3) can be summarized as follows: 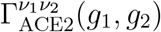 is the number of conformations in which two glycans overlap (any given conformation counts as a 0 or 1), whereas the density overlap (r.h.s. of Eq. (3)) depends on the number of clashes, which can be larger than 1 for a given conformation of the two glycans in which they clash. Equation (3) is instructive because the calculation of the glycan density requires only the exhaustive exploration of single-chain conformations.

In Table 1 we show the result of the upper bound for all pairs of ACE2 sites that could be glycosylated, with the same glycan attached to all the sites. The diagonal entries are just a reference, reflecting the unphysical interaction of an ensemble of glycans with itself. Most of the off-diagonal entries are small or zero, indicating that the independence approximation for the corresponding pair of glycans is excellent. The only exception is the bound for the oligosaccharides at sites N322-N546. In this case, there is a large overlap in the conformational space, which results in the reduction of the volume available for both of these carbohydrates. Although predicting a priori how this repulsion modulates the affinity for RBD is complicated, we can use the structure of ACE2 to suggest a hypothesis. The N546 glycan is further away from the RBD-binding interface than the N322 oligosaccharide. As a consequence, (i) the N546 glycan does not directly impact RBD binding (Fig. 4), and (ii) we expect that it reduces only (or predominantly) the number of conformations of the N322 oligosaccharide in which this glycan is located away from the RBD. Under these premises, the presence of the N546 glycan favors N322 oligosaccharide conformations that explore the RBD-binding interface, thereby enhancing the shielding effect of the N322 carbohydrate and resulting in an increase of the dissociation constant. Although plausible, our hypothesis cannot be rigorously proved, but it can be experimentally tested by measuring the dissociation constant of ACE2 constructs that are incapable of N-linked glycosylation at N546. Overall, we surmise that neglecting the correlation between glycans bound at two residues contributes little to the dissociation constant.

**Table 1:**
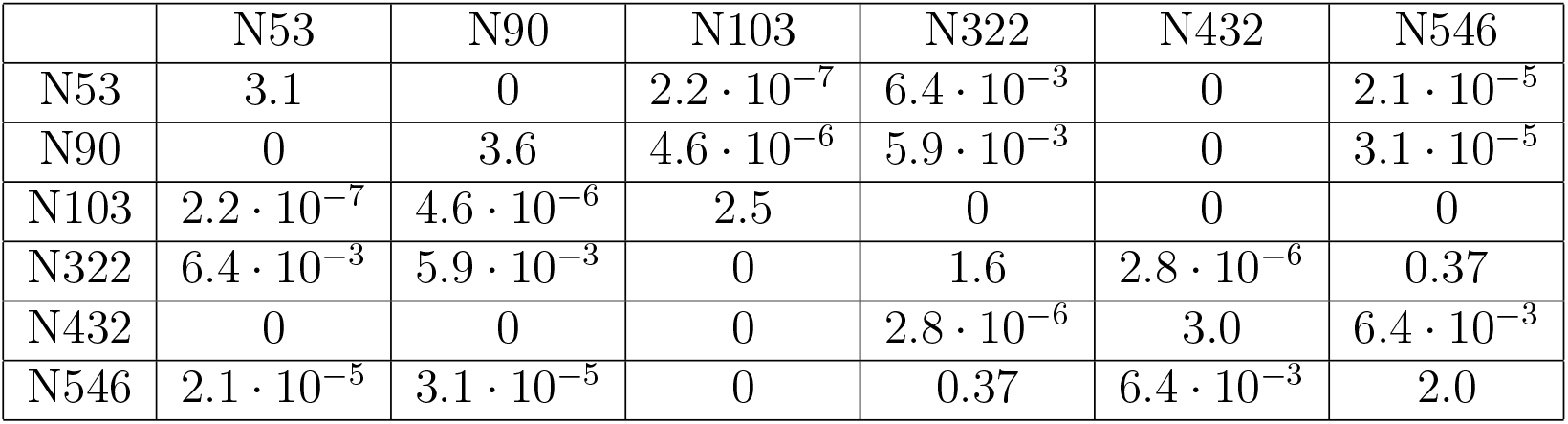
Upper bound in the r.h.s. of Eq. 3. The same glycan is attached to all sites. The numbers refer to glycan A3G3S11, with degree of polymerization 13, for *a* = 5.5 Å. The attachment site for a glycan was chosen as the closest site to the nitrogen of the carboxyamine of the the corresponding asparagine. The vertex corresponding to N103 was completely enclosed by occupied sites, therefore we attached the oligosaccharide to a nearby vertex.

Although our theory describes the experimental data nearly quantitatively, we need to make a few comments to put it in context in order to discern the discrepancy between experiments and theory. (1) The statistical error in *K*_*D*_*/K*_*D*_(∅) for a few experiments is absent or underestimated because the error bar for either *K*_*D*_(∅) or both *K*_*D*_ and *K*_*D*_(∅) was not reported. (2) The *p*_*ν*_(*g*)’s are not exact. Estimating the errors in their values is challenging. For this reason, we took two sets of *p*_*ν*_(*g*) obtained from different groups^17,18^ to gauge what would be the impact of inaccurate estimate of these probabilities. In at least a few cases, the difference in the results of the model obtained using the two sets of *p*_*ν*_(*g*) is on the order of the discrepancy from experiments. (3) The inaccurate recovery of the effect of the ablation of the N90 glycan might occur if T92Q changes the affinity for RBD regardless of the glycan interaction. As shown in the right panel of Fig. 3, the model compares more favorably with results obtained by adopting a different modification of ACE2 (T92I, see ref.^31^) that prevents the attachment of the glycan at N90 (compare with the triangle pointing right). The impact of the mutation could be tested by measuring the dissociation constant of T92Q and T92I after ablating all the glycans with PNGase F. (4) Obtaining much higher accuracy in comparison with experiments presents some challenges. Experimental measurements for the dissociation constant from various groups change within three orders of magnitude (see Fig. S3). Even if we consider only the WT results, the range goes from *K*_*D*_ ≈ 1 nM to *K*_*D*_ ≈ 100 nM (see Fig. S3). Differences in experimental setup contribute to this large variability. For instance, not all of the studies used the same segment of the spike protein (some use the whole S-protein, ^18,30^ others only the RBD^25,26,31^). In addition, some studies were performed with monomeric ACE2 (e.g. with his-tag^18,31^), while others performed measurements using a dimerized form of the receptor (e.g. with Fc-tag ^26,31^).

Even after considering all these factors, we conclude that it is remarkable that after scaling the data by the dissociation constant for the WT or deglycosylated ACE2, the results fall within a factor of 2 to 3 in most cases, which hints at a simple and general mechanism that is independent of the variabilities in the experimental setups. Our model provides this rationale: entropic effects dominate over specific interactions, whose contribution might at most account for changes on the order of 1 *k*_*B*_*T* or less, under the strong assumption that the RBD-glycan energy is the same for all the conformations of the glycans.

## Conclusions

In conclusion, we have provided numerical evidence showing that ACE2 glycans repel RBD binding by weak entropic forces. This prediction may be tested experimentally by measuring the temperature dependence of the ratio between dissociation constants of WT and deglycosylated ACE2. If our prediction holds, changing temperature (not relevant biologically perhaps) over a limited range should not affect the results significantly. In addition, we have predicted the strength of the impact on ACE2-RBD binding of glycans attached to various ACE2 sites (Fig. 4). We have argued that the N546 oligosaccharide plays an indirect role in modulating ACE2-RBD binding.

A consequence of our theory and the experimental results that we used to validate the predictions is that the spike protein of SARS-CoV-2 does not specifically recognize the ACE2 glycans. However, carbohydrates are known to impact the function of SARS-CoV-2 virus. For instance, complete deglycosylation of ACE2 strongly impacts viral entry, possibly because it affects the expression of the receptor on host cell surface.^28^ In addition, other carbohydrates in the host glycocalyx seem to aid SARS-CoV-2 binding, in particular heparan sulfate^43^ and glycolipids rich in sialic acids.^44^ Moreover, glycan remodeling indicated that neutral forms of the N322 RBD glycan strengthen the interaction with ACE2.^32^

Finally, we extend our analysis to the interaction between the spike protein and the immune system, under the assumption that antibodies do not specifically recognize the glycans belonging to the spike.^45^ Let there be *N*_*g*_ = 18 glycosylation sites on the spike, and let *R*_*g*_ ≈ 7.5 Å be the radius of glycans (from our MD calculations); let *R*_*a*_ ≈ 18.6 Å be the size of an antibody (see^12^), and let *A* ≈ 84255 Å^2^ be the antibody accessible surface area of the protein.^12^ If an antibody interacts with at most one glycan of the spike, the percentage of spike protein shielded is given by 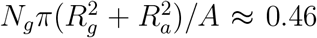, which is in good agreement with a more sophisticated analysis by Woods and coworkers (≈ 40%, see^12^). Therefore, to a first approximation, antibodies should interact with at most one glycan at a time. If this is the case, our model suggests that the glycan shield should have a small effect on antibody binding, with the exception of small areas of the spike surface, where putative epitopes would overlap with glycosylation sites leading to *K*_*D*_*/K*_*D*_(∅) ≈ Ω_Spike_*/*Ω_Spike·Antibody_ ≫ 1.

## Conflict of Interest

There are no conflicts to declare.

## Acknowledgement

We thank Hung Nguyen for useful discussions. This work was supported by a grant from the National Science Foundation (CHE19-000033) and the Welch Foundation through the Collie-Welch Chair (F-0019).

## Notes

### Competing Interest Statement

The authors have declared no competing interest.

