## Supplementary Information for "Reduction in RBD Binding Affinity to Glycosylated ACE2 is Entropic in Origin"

¶*Current Address: Institute for Soft Matter Synthesis and Metrology, Georgetown University, Washington, DC 20057*

#### Derivation of Eqs. 1-2

**Intuitive Argument:** Let the dissociation constant of the complex between RBD and a N-glycosylated ACE2 be,

$$K_D(\{g_\nu\}) = e^{\beta \Delta F^0(\{g_\nu\})}, \tag{S1}$$

where $\Delta F^0(\{g_\nu\})$ is the change in the binding free energy between the bound (ACE2·RBD) and unbound states. Note that with the symbol $\{g_\nu\}$ we indicate concisely the dependence on all of the 6 N-linked glycans, that is $\Delta F^0(\{g_\nu\}) = \Delta F^0(g_{\mathrm{N53}}, g_{\mathrm{N90}}, g_{\mathrm{N103}}, g_{\mathrm{N322}}, g_{\mathrm{N432}}, g_{\mathrm{N546}})$. Here, $g_\nu$ is an index that indicates which glycan is linked to asparagine $\nu$, with $\nu \in \mathcal{N} = \{\mathrm{N53}, \mathrm{N90}, \mathrm{N103}, \mathrm{N322}, \mathrm{N432}, \mathrm{N546}\}$. We ignore any contribution from RBD glycans.

We make a series of approximations to simplify the formally exact relation in Eq S1. (i) We assume that the internal degrees of freedom of the two proteins (ACE2 and RBD) are frozen. (ii) Similarly, we neglect the relative movement between ACE2 and the RBD in the bound state. In practice, for approximations (i) and (ii) to hold, it is sufficient that the fluctuations of the proteins are small enough that they do not change the mapping to the lattice of ACE2 and the ACE2-RBD complex. In this way, proteins (ACE2 and RBD) and ACE2 glycans are decoupled, the partition function is separable into product of a term neglecting the glycans and featuring only protein degrees of freedom, and another term accounting for glycan degrees of freedom while the proteins are "frozen". (iii) We approximate the term involving only the proteins as the free energy of the de-glycosylated ACE2 by itself or bound to the RBD. (iv) Finally, we assume that the glycans interact with the protein and with other polysaccharides via self-avoiding interactions, which means that their energy is irrelevant. Thus, the glycan contribution to the free energy of binding is entropic.

Under these premises, we write Eq. S1 as:

$$\begin{aligned} K_D(\{g_\nu\}) &= e^{\beta \Delta F^0(\emptyset)} e^{k_{\mathrm{B}}^{-1}[S^0_{\mathrm{ACE2}}(\{g_\nu\}) - S^0_{\mathrm{ACE2 \cdot RBD}}(\{g_\nu\}]} = \\ &= K_D(\emptyset) e^{k_{\mathrm{B}}^{-1}[S^0_{\mathrm{ACE2}}(\{g_\nu\}) - S^0_{\mathrm{ACE2 \cdot RBD}}(\{g_\nu\})]} \end{aligned} \tag{S2}$$

where $e^{\beta \Delta F^0(\emptyset)} = K_D(\emptyset)$ is the dissociation constant in the absence of glycans, and $S^0_{\mathrm{ACE2}}(\{g_\nu\}) - S^0_{\mathrm{ACE2 \cdot RBD}}(\{g_\nu\})$ is the difference in ACE2 N-linked glycan entropy between the unbound and bound states. Next, we replace the entropy with the logarithm of the number of conformations ($\Omega_{\mathrm{ACE2}}$ and $\Omega_{\mathrm{ACE2 \cdot RBD}}$ for the unbound and unbound cases, respectively) and obtain,

$$K_D(\{g_\nu\}) = K_D(\emptyset) \frac{\Omega_{\mathrm{ACE2}}(\{g_\nu\})}{\Omega_{\mathrm{ACE2 \cdot RBD}}(\{g_\nu\})}. \tag{S3}$$

The number of conformations, $\Omega_{\mathrm{ACE2}}$ and $\Omega_{\mathrm{ACE2 \cdot RBD}}$, are impossible to evaluate exhaustively. This is because there are 6 polysaccharides bound to ACE2, and on a 3-dimensional cubic lattice each one of them has about $\omega \approx 10^3 - 10^9$ conformations (see Table S1). Even if only two of them could occupy each other's space, the number of conformations to compute would be $\Omega \approx \omega^2 \approx 10^6 - 10^{18}$, which makes the enumeration of all self-avoiding conformations extremely challenging if not impossible (this is also the reason why we ignore the interaction between glycans of the spike protein and ACE2 glycans in this model). In order to make the calculations manageable, we assume that the N-linked glycans of ACE2 are independent, that is, they do not interact with each other, not even by volume exclusion. Therefore, we can write the number of glycan conformations as the product of the number of conformations allowed for each polysaccharide in the absence of the others, which results in,

$$K_D(\{g_\nu\}) = K_D(\emptyset) \prod_{\nu \in \mathcal{N}} \frac{\Omega_{\mathrm{ACE2}}(g_\nu)}{\Omega_{\mathrm{ACE2 \cdot RBD}}(g_\nu)}. \tag{S4}$$

This is Eq. 1 of the main text.

In order to account for the heterogeneity of ACE2 glycoforms, we average over $p(\{g_\nu\})$, which is the probability that a specific glycoform $\{g_\nu\}$ occurs. That is,

$$\overline{K_D} = K_D(\emptyset) \sum_{g_{\mathrm{N53}}} \cdots \sum_{g_{\mathrm{N546}}} p(g_{\mathrm{N53}}, \cdots, g_{\mathrm{N546}}) \prod_{\nu \in \mathcal{N}} \frac{\Omega_{\mathrm{ACE2}}(g_\nu)}{\Omega_{\mathrm{ACE2 \cdot RBD}}(g_\nu)}. \tag{S5}$$

Under the assumption that there is no correlation between the glycans N-linked to two different asparagines, we can write $p(g_{\mathrm{N53}}, \cdots, g_{\mathrm{N546}}) = \prod_{\nu \in \mathcal{N}} p_\nu(g_\nu)$, where $p_\nu(g)$ denotes a probability of finding glycan $g$ N-linked to asparagine $\nu$. Using this last expression, we finally obtain:

$$\overline{K_D} = K_D(\emptyset) \sum_{g_{\mathrm{N53}}} \cdots \sum_{g_{\mathrm{N546}}} \prod_{\nu \in \mathcal{N}} p_\nu(g_\nu) \prod_{\nu \in \mathcal{N}} \frac{\Omega_{\mathrm{ACE2}}(g_\nu)}{\Omega_{\mathrm{ACE2 \cdot RBD}}(g_\nu)}, \tag{S6}$$

and by re-ordering products and summations we obtain,

$$\overline{K_D} = K_D(\emptyset) \prod_{\nu \in \mathcal{N}} \Big[ \sum_{g} p_\nu(g) \frac{\Omega_{\mathrm{ACE2}}(g)}{\Omega_{\mathrm{ACE2 \cdot RBD}}(g)} \Big], \tag{S7}$$

which is Eq. 2 in the main text.

Note that the average over the glycoforms introduced in Eq. S5 is annealed. This choice is justified if experiments report on the average of the dissociation constant, and not the binding free energy, over an ensemble of different glycoforms, that is $K_D^{\rm exp} = K_D$. If the experiments were to measure the free energy averaged over the glycoforms, then we should have performed a quenched average, that is $K_D^{\rm exp} = e^{\beta \overline{\Delta F^0}}$. Strictly speaking, the output of Surface Plasmon Resonance experiments is the off- and on-rate of RBD binding to ACE2 averaged over multiple events likely involving different glycoforms of ACE2, thus $\overline{k_{off}}$ and $\overline{k_{on}}$. As a result the dissociation constant is, $K_D^{\rm exp} = \overline{k_{off}}/\overline{k_{on}}$. This is clearly close to, although not exactly the same as the annealed average in Eq. S5.

### Sampling the Conformations

Glycans are modeled as branched polymers on a 3-dimensional cubic lattice. Each sugar in the polysaccharide occupies one vertex, and all the self-avoiding conformations are allowed. In Fig. S1, we illustrate the initial steps of the algorithm used to sample exhaustively the conformations of the glycans. First, we define an anchor site (black sphere, numbered 0) which is fixed and is used to place the polysaccharide chain on a lattice site occupied by the side chain of an ACE2 asparagine. Next, we add the first sugar (an N-acetyl-glucosamine, or GlcNAc, shown as a blue cube numbered 1). Rotations around the anchor place bead 1 on 6 vertices. The choice among these rotations will be discussed later. For now the position of the first bead is fixed. Then, we add the second sugar (again, GlcNAc, shown as a blue cube numbered 2), which is attached to bead 1. Next, we add the first mannose (green sphere, numbered 3) to bead 2. The resulting chain is linear and has degree of polymerization 3 (the anchor is not included in the degree of polymerization). Beads numbered 4 and 5, two mannoses (green spheres), are added next. This completes the shared core of all N-linked glycans. The assembly of more complex polysaccharides proceeds following the same process of adding the sugars one-by-one. All conformations are allowed, except if two beads occupy the same lattice site.

The list of all the glycans considered in this study as a function of the degree of polymerization is shown in Fig. S2, and the number of conformations for the free glycan are reported in Table S1. The reader might have observed by now that certain conformations of the sugar are identical up to permutations of the labels (see the fifth and sixth structures in the box at the bottom of Fig. S1). We did not prune the ensemble of conformations to remove these identical copies related by permutations of the indices. The reasons are that (i) in real glycans, sugar 4 and 5 (both of them mannose) differ. For instance, they are attached to different sites of the sugar ring of the second GlcNAc. In addition, (ii) the final result of the paper (i.e. the impact of glycan entropy to the dissociation constant) would be unaffected by removing the identical conformations. This is because we only care about the ratio of the number of allowed conformations with and without the RBD. If we do not prune the ensemble both the numerator and the denominator are overestimated by the same multiplicative factor, which cancels out.

### Conformations of the Glycan linked to ACE2

For each asparagine $\nu$ at the beginning of the N-X-S/T sequon of ACE2, we construct a discretized version of ACE2 centered on the nitrogen of the asparagine side-chain amide group. We proceed by constructing the set of vertices, which are occupied by the protein. In order to do so, we consider a lattice site to be occupied if the minimum distance, $r_{min}$, between this vertex all heavy atoms on the protein satisfies $r_{min} \leq a/2 + r_{\text{ha}}$, where $a = 0.55$ nm is the lattice spacing, which is roughly commensurate with the distance between the centers of mass of two consecutive sugars in a polysaccharide chain (see Fig. 2A), and $r_{\text{ha}}$ is the van der Waals radius of a protein heavy atom[S1] (1.79 Å, 1.55 Å, 1.52 Å, and 1.80 Å for C, N, O, and S, respectively). Next, we place the anchor of a glycan (see Fig. S1) at the origin of the cubic lattice (the location of the side-chain of the chosen asparagine) and we identify the direction of the link connecting the anchor and the first sugar which results in conformations that do not clash with the protein. For each one of these directions, we run through the ensemble of structures for each polysaccharide, and we count all the self-avoiding conformations in the presence of ACE2. We repeat the operation in the presence of the RBD.

#### Calculation of Glycoform probabilities

The probabilities $p_\nu(g)$ in Tables S2, S4, S6 are taken from Supplementary Tables 1-2 of Crispin and coworkers.[S2] Some manipulation of the data was necessary in order to obtain the probabilities. First, in this work we ignore fucosylation and hybrid glycans. Therefore, the data for glycans FA1 and A1, FA2 and A2 etc are combined. Second, we ignore the possibility of bisecting glycans. Therefore, as an example, the data for A2/A1B is fully assigned to A2. Third, the experimental data is apportioned between agalactosylated, galactosylated, and galactosylated and sialylated glycans, without specifying the number of galactose or sialic acid moieties. Thus, we split the probability evenly among all of the constructs that we have considered that match this description. For instance, galactosylated and not sialylated A3/A2B and FA3/FA2B corresponds to A3G10, A3G01, A3G11, A3G02, and A3G3. In the case of WT N53, this moiety appears about $\approx (15 - 16)\%$ of the time, which means that each one of the 5 corresponding polysaccharides in this study is given a probability of 0.031. In order to construct the probabilities $p_\nu(g)$ for the case in which sialidase is added to WT or +ST6 ACE2 enzymes (Tables S3, S5), we assume that all of the sialic acids have been removed. This means that we assign the probability of a sugar ending with a sialic acid to the corresponding galactosilated but not sialylated glycan. For example, in the case of N53 for the +ST6 ACE2, A3/A2B and FA3/FA2B are $\approx 1\%$ galactosylated, and $\approx 73\%$ sialylated. Thus, for the +ST6+Sia experiment with the same glycan, we assign 74% to galactosylated A3/A2B and FA3/FA2B. Next, as explained before, we assign to all of the glycans considered in this study that match the description (A3G10, A3G01, A3G11, A3G02, and A3G3) the same probability (0.147).

The probabilities in Table S7 are extracted using WebPlotDigitizer[S3] from Fig. 4A by Azadi and coworkers.[S4] Again, we performed some minimal manipulations. First, we ignored fucosylation, thus we combined the probabilities for glycans distinguished only by the presence of fucose. Second, if there is ambiguity in terms of the location of a sugar ring, we assigned an equal probability to all of the glycans that match the description from Fig. 4A by Azadi and coworkers.[S4] Finally, for N322 Azadi and coworkers report a polysaccharide larger than any of the ones considered here. We ignored that, which is likely a negligible simplification given that its probability is very small. When constructing the probabilities $p_\nu(g)$ after sialidase is added to solution, we followed the same procedure outlined before: we take the WT data and combine and assign the populations of polysaccharides ending with a sialic acid to the closest glycan ending with a galactose.

### Impact of the lattice spacing value

We tested the impact of changing the lattice spacing value from $a = 5.0$ Å to $a = 6.0$ Å. As shown in Fig. S4, qualitatively the results for the dissociation constant are not affected by changing the lattice spacing. On the other hand, the dependence of the contribution of each asparagine amenable to glycosylation on the degree of polymerization of the oligosaccharides is affected by the resolution of the lattice. However, the general trend is consistent, which indicates that the precise value of the **only free parameter** in the lattice model does not drastically impact the results.

### Proof of Eq. 3

Consider a pair of ACE2 sites, $\nu_1$ and $\nu_2$, that are amenable to N-glycosylation. If these two are treated as independent, there are $\Omega^{\nu_1}_{\mathrm{ACE2}}(g_1)\Omega^{\nu_2}_{\mathrm{ACE2}}(g_2)$ oligosaccharide conformations, where $g_1$ and $g_2$ are the glycans N-linked to $\nu_1$ and $\nu_2$. Let $\Gamma^{\nu_1\nu_2}_{\mathrm{ACE2}}(g_1, g_2)$ be the number of conformations in which the glycans $g_1$ and $g_2$ clash on one or more lattice sites. Then, the number of self-avoiding conformations is given by,

$$\Omega^{\nu_1}_{\text{ACE2}}(g_1)\Omega^{\nu_2}_{\text{ACE2}}(g_2) - \Gamma^{\nu_1\nu_2}_{\text{ACE2}}(g_1,g_2) = \Omega^{\nu_1}_{\text{ACE2}}(g_1)\Omega^{\nu_2}_{\text{ACE2}}(g_2)\Big[1 - \frac{\Gamma^{\nu_1\nu_2}_{\text{ACE2}}(g_1,g_2)}{\Omega^{\nu_1}_{\text{ACE2}}(g_1)\Omega^{\nu_2}_{\text{ACE2}}(g_2)}\Big]. \tag{S8}$$

As long as the ratio in the squared bracket on the r.h.s. is small, we can ignore the correction, and the independence approximation would be fairly accurate. To test whether this is the case without explicitly computing $\Gamma^{\nu_1\nu_2}_{\text{ACE2}}(g_1,g_2)$ (a daunting task), we searched for an upper bound for the ratio $\Gamma^{\nu_1\nu_2}_{\text{ACE2}}(g_1,g_2)/[\Omega^{\nu_1}_{\text{ACE2}}(g_1)\Omega^{\nu_2}_{\text{ACE2}}(g_2)]$. Before deriving the bound, let us simplify the notation by defining $\Gamma_{12} = \Gamma^{\nu_1\nu_2}_{\text{ACE2}}(g_1,g_2)$, $\Omega_1 = \Omega^{\nu_1}_{\text{ACE2}}(g_1)$, and $\Omega_2 = \Omega^{\nu_1}_{\text{ACE2}}(g_2)$. We can now define,

$$\frac{\Gamma_{12}}{\Omega_1\Omega_2} = \frac{1}{\Omega_1\Omega_2}\sum_{c_1\in\Omega_1}\sum_{c_2\in\Omega_2}\Theta\Big(\sum_{\alpha\in c_1}\sum_{\beta\in c_2}\delta_{\alpha,c_1;\beta,c_2}\Big). \tag{S9}$$

Here, $c_1$ ($c_2$) is the index of conformations in the exhaustive ensemble $\Omega_1$ ($\Omega_2$), whereas the index $\alpha$ ($\beta$) runs over the lattice vertices occupied by the glycan in conformation $c_1$ ($c_2$). The step function $\Theta$ is equal to 1 if the argument is $> 0$, and is equal to zero otherwise, and the Kronecker delta, $\delta_{\alpha,c_1;\beta,c_2}$, is equal to 1 if vertex $\alpha$ from conformation $c_1$ of the first glycan is the same as vertex $\beta$ of conformation $c_2$ of the second glycan, otherwise it is zero. What this means is that we sum over all the conformations of the two glycans, and we count the number of pairs of conformations (one for the first oligosaccharide, another for the second one) for which there is at least one clash. A very simple bound of this equation can be obtained as follows:

$$\frac{\Gamma_{12}}{\Omega_1\Omega_2} \le \frac{1}{\Omega_1\Omega_2}\sum_{c_1\in\Omega_1}\sum_{c_2\in\Omega_2}\sum_{\alpha\in c_1}\sum_{\beta\in c_2}\delta_{\alpha,c_1;\beta,c_2}, \tag{S10}$$

which is obtained from Eq. S9 by bringing out the argument of the step function $\Theta$. To understand why this is an upper bound, let us make a few examples. (i) If there are no clashes between conformation $c_1$ of the first glycan and conformation $c_2$ of the second one, then $\sum_{\alpha\in c_1}\sum_{\beta\in c_2}\delta_{\alpha,c_1;\beta,c_2} = 0$, and $\Theta(\sum_{\alpha\in c_1}\sum_{\beta\in c_2}\delta_{\alpha,c_1;\beta,c_2}) = 0$. Thus, the contribution of this pair of conformations to Eq. S9 and Eq. S10 is the same. (ii) If there is only one clash, that is only one sugar of the first glycan in conformation $c_1$ occupies the same lattice site as a sugar of the second glycan in conformation $c_2$, then $\sum_{\alpha\in c_1}\sum_{\beta\in c_2}\delta_{\alpha,c_1;\beta,c_2} = 1$ and $\Theta(\sum_{\alpha\in c_1}\sum_{\beta\in c_2}\delta_{\alpha,c_1;\beta,c_2}) = 1$. Therefore, also in this case the contribution to Eq. S9 and Eq. S10 of this pair of conformations is identical. (iii) Let there now be multiple clashes between the first glycan in conformation $c_1$ and the second in conformation $c_2$, that is the two polysaccharides share multiple lattice sites. In this case, $\sum_{\alpha\in c_1}\sum_{\beta\in c_2}\delta_{\alpha,c_1;\beta,c_2} > 1$, but because of the $\Theta$ function, $\Theta(\sum_{\alpha\in c_1}\sum_{\beta\in c_2}\delta_{\alpha,c_1;\beta,c_2}) = 1$. As a consequence, this pair of conformations featuring multiple clashes contributes with a larger value to Eq. S10 than to Eq. S9. To sum up, the numerator of the r.h.s. of Eq. S10 counts the number of clashes between $g_1$ and $g_2$ in their conformational ensembles, whereas the numerator in Eq. S9 count the number of pairs of conformations in which there is at least one clash, which is smaller than the total number of clashes.

Next, we use the following identity:

$$\delta_{\alpha\beta} = \sum_{i\in\text{lattice}} \delta_{\alpha i}\delta_{i\beta},$$

where the index $i$ runs over all the vertices in the cubic lattice. We plug this expression in Eq. S10 and after minor manipulations we obtain,

$$\begin{aligned}\frac{\Gamma_{12}}{\Omega_1\Omega_2} \leq & a^6 \sum_{i\in\text{lattice}} \left\{ \left[ \frac{1}{a^3\Omega_1} \sum_{c_1\in\Omega_1}\sum_{\alpha\in c_1} \delta_{\alpha i} \right] \left[ \frac{1}{a^3\Omega_2} \sum_{c_2\in\Omega_2}\sum_{\beta\in c_2} \delta_{i\beta} \right] \right\} = \\ & a^6 \sum_{i\in\text{lattice}} \rho_1(i)\rho_2(i)\end{aligned} \tag{S11}$$

Here, the expressions in squared brackets are the densities ($\rho$) of the first and the second glycan at lattice site $i$, averaged over the ensemble of glycan conformations. This proves Eq. 3 in the main text.

We conclude this section noting the similarity between the bound in Eq. S11 and the argument proposed by Flory to obtain the exponent $\nu$ indicating the power law dependence of the size of a linear polymer ($R_g$) on the length of the polymer ($N$), $R_g \sim N^{\nu}$.[S5] Flory minimized a free energy including an entropic term derived from the ideal chain, and an interaction term featuring a term similar to the r.h.s. of Eq. S11. As discussed by De Gennes,[S5] these two terms overestimate the exact values. However, they balance out, yielding a nearly exact value of $\nu$.

### MD Simulations in Explicit Water

We performed all-atom MD simulations of glycans (PDB ID: 5GSQ[S6]) in explicit solvent (water) based on the CHARMM36 force field.[S7] We considered three different kinds of simulation setup: a single glycan, a pair of glycans, and four chains of glycans in a periodically replicated cubic box. For all these systems, the glycans were solvated with TIP3P water[S8] and 150mM KCl using Visual Molecular Dynamics.[S9] After 2,000 iterations of energy minimization, each system was equilibrated for 1 ns with the time step of 2 fs prior to a production run, at 1 bar and 300K using a Nose-Hoover thermostat[S10] and a Parrinello-Rahman barostat.[S11] The long-range part of electrostatic interactions was handled by the Particle Mesh Ewald method.[S12] All energy minimizations and simulations were performed using LAMMPS.[S13]

For a single glycan, we chose 48 Å for the width of the simulation box. Since the end-to-end distance for the glycan's most stretched conformation is only about 20 Å, the influence of the periodic images is negligible. We ran the simulation for 200 ns from which the configuration was sampled every 2 ps for the statistics.

For a pair of glycans, we apply a biasing potential, $u_{\text{bias}}(r; r_0) = \frac{1}{2}k(r - r_0)^2$, which represents a harmonic spring rendering the pair distance $r$ between the centers of mass of the two glycans to fluctuate around $r_0$. We used $k = 1.8\,\text{kcal}/(\text{mol} \cdot {}^2)$ for the spring constant and prepared the initial configurations with the various pair distance ranging from $r_0 = 5$ to $r_0 = 35$ with the increment of $\Delta r_0 = 1$. The width of the simulation box was 60 Å, and thus, for $30 \leq r_0 \leq 35$, the glycan pair was arranged to be along the diagonal of the cubic box.

For each system with a given $r_0$, we ran the simulation for 50 ns from which the configuration was sampled every 0.2 ps. To ensure that the production run is long enough, we computed the time correlation function for the relative orientation between two glycans, using,

$$C(t) = \left\langle \left[\hat{\mathbf{R}}^{(1)}_{\text{ee}}(t) \cdot \hat{\mathbf{R}}^{(2)}_{\text{ee}}(t)\right] \left[\hat{\mathbf{R}}^{(1)}_{\text{ee}}(0) \cdot \hat{\mathbf{R}}^{(2)}_{\text{ee}}(0)\right] \right\rangle , \tag{S12}$$

where $\hat{\mathbf{R}}^{(1)}_{\text{ee}}(t)$ and $\hat{\mathbf{R}}^{(2)}_{\text{ee}}(t)$ are the end-to-end unit vectors of each glycan at time $t$. The correlation function decays faster as $r_0$ increases (Fig. S7A). The correlation time, $\tau$, estimated from $C(\tau)/C(0) = 1/e$, is less than 5 ns for all $r_0$ except $r_0 = 5$, which gives $\tau \approx 19$ ns as the glycan pair is pulled against each other at too close a distance (Fig. S7B). Therefore, we can regard the sampled pair configurations for $r_0 \geq 6$ to be in equilibrium. From the obtained statistics, we construct the free energy curve (PMF) using the standard Weighted Histogram Analysis Methods (WHAM).[S14]

For the 4-glycan system, we prepared the initial configurations in which glycans are equidistant from each other with the pair distance of $\sqrt{L}/2$ where $L$ is the box width. We simulated the systems with $L$ = 48 Å, 58 Å, 73 Å, 88 Å, and 104 Å. The production run was 200-ns long for $L = 48$ Å and 58 Å, and was longer than 200 ns for the larger systems. During the production runs, the configuration was sampled every 10 ps. To make sure that the simulation time is long enough, we calculated an ergodic measure[S15–S17] that quantifies the deviation between each individual pair distance, $d_{i,j}(t)$, and the distance averaged over all the pairs, $d(t) = \frac{1}{6}\sum_{i<j} d_{i,j}(t)$. The ergodic measure for a given trajectory is defined as,

$$\omega(t) = \frac{1}{6L^2} \sum_{i=1}^{3} \sum_{j=i+1}^{4} |\bar{d}_{i,j}(t) - \bar{d}(t)|^2 , \tag{S13}$$

where $\bar{d}_{i,j}(t)$ and $\bar{d}(t)$ are the time averages along the trajectory, given by $\bar{d}_{i,j}(t) = \frac{1}{t}\int_0^t d_{i,j}(t')\mathrm{d}t'$ and $\bar{d}(t) = \frac{1}{t}\int_0^t d(t')\mathrm{d}t'$, respectively. If the system is ergodic, $\omega(t)$ should approach zero asymptotically. In Fig. S8E, $\omega(t)$ decays below 0.001 within 200 ns for $L = 48\,\text{Å}$ and 58 Åwhereas the decay is relatively slower for the larger systems. It is notable that the trajectory for $L = 88\,\text{Å}$ shows a particularly slow decay of $\omega(t)$ due to the rare fluctuations at an earlier time in the trajectory, resulting in more frequent contacts than expected (Fig. S8C).

Figure S8A shows the distribution, $P(r/L)$, of the distance between two glycans for various box sizes, $L$. We observe that the distributions for different box size collapse onto one master curve except for $L = 48$ Å. The shape of the curve is a consequence of the periodic boundary conditions and the minimum image convention, which result in the distances between two particles along a Cartesian axis to be $\leq L/2$. Therefore, if a particle is at the center of the periodic box, the image of the other particles to be considered is inside a box of side $L$. For an ideal gas, the probability density that two particles are at distance $r$ is proportional to the surface of a sphere of radius $r$, that is $P(r) \propto r^2$ (black dashed line in Fig. S8A). As long as $r < L/2$, the whole sphere is inside the box of side $L$, so the whole sphere is to be considered and $P(r)$ is indeed quadratic in $r$. For $L/2 \leq r < \sqrt{3}/2L$ part of the sphere is outside the box and should not be computed, so $P(r)$ decreases with $r$ until $r = \sqrt{3}/2L$, after which it must be zero. This shape is roughly recapitulated in Fig. S8A, where the curves grow up to $\approx L/2$ and then decrease. What matters here is that for $r < L/2$ (the only domain that is relevant), $P(r) \approx 0$ for $r \lessapprox 5$ Å, indicating a strong repulsion between the glycans. Next there is a rapid rise, which suggests some "stickiness" for distances between $5\,\text{Å} < r < 15\,\text{Å}$. After that, $P(r)$ looks almost quadratic in $r$ up to about $r = L/2$, and then decreases. The general pictured that emerges from this analysis is in rough agreement with the PMF in Fig. 2C of the main text.

The contact probability was computed using $P_{\text{cont}} = \langle \Theta(R_c - d_{i,j}) \rangle$, where $R_c$ is the cutoff distance for a contact and $\Theta(x)$ is the Heaviside step function ($\Theta(x) = 1$ if $x > 0$ and $\Theta(x) = 0$ if $x < 0$). We used $R_c = 15\ \approx 2\langle R_g \rangle$, where $\langle R_g \rangle$ is the mean of the radius of gyration obtained from the single-glycan simulation. Nevertheless, if we compute the aggregation probability that more than two glycans cluster with one another, using

$$P_{\mathrm{aggr}} = \left\langle \min \left[ \sum_{i \neq j \neq k}^{4} \Theta(R_c - d_{i,j}) \Theta(R_c - d_{i,k}), 1 \right] \right\rangle \ , \tag{S14}$$

For an ideal gas, one expects that the contact probability decreases as the volume of the box, so $P_{\mathrm{cont}} \propto L^{-3}$. Similarly, the probability of forming clusters of 3 glycans should be inversely proportional to the square of the volume of the box, that is $P_{\mathrm{aggr}} \propto L^{-6}$. As shown in Fig. S8D, these relationships are nearly quantitatively captured in our simulations, indicating the lack of stable dimers or trimers during the timescale of our simulations.

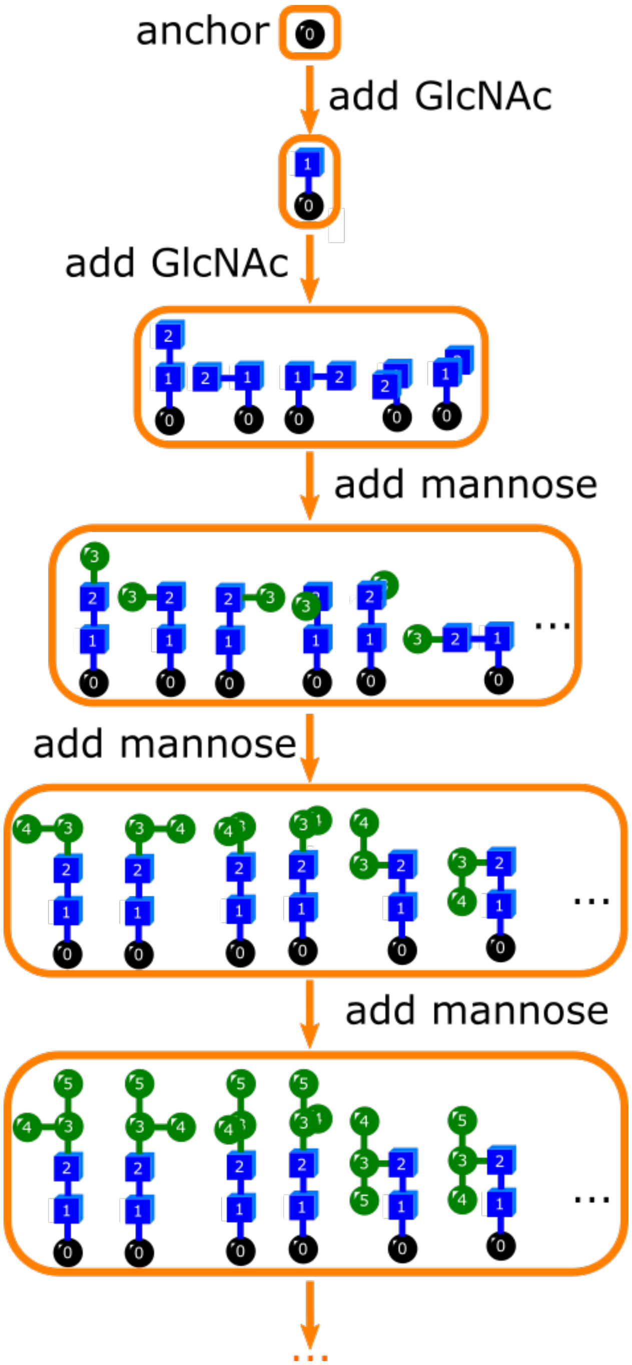


Figure S1: Procedure for exhaustive sampling of the glycan conformations. First (top orange box), set the anchor (black sphere, number 0), which corresponds to the position occupied by the side chain of a given ACE2 asparagine. Then, add the first bead (GlcNAc, blue square, number 1). This bead is assigned a fixed orientation at the beginning. Next, add the second bead (GlcNAc, blue square, number 2) to bead number 1. Subsequently, insert bead 3 (green, mannose) to bead 2. Finally, introduce the fourth and fifth beads (green, mannose) to bead number 3. The algorithm continues until the full topology of the glycan is constructed.

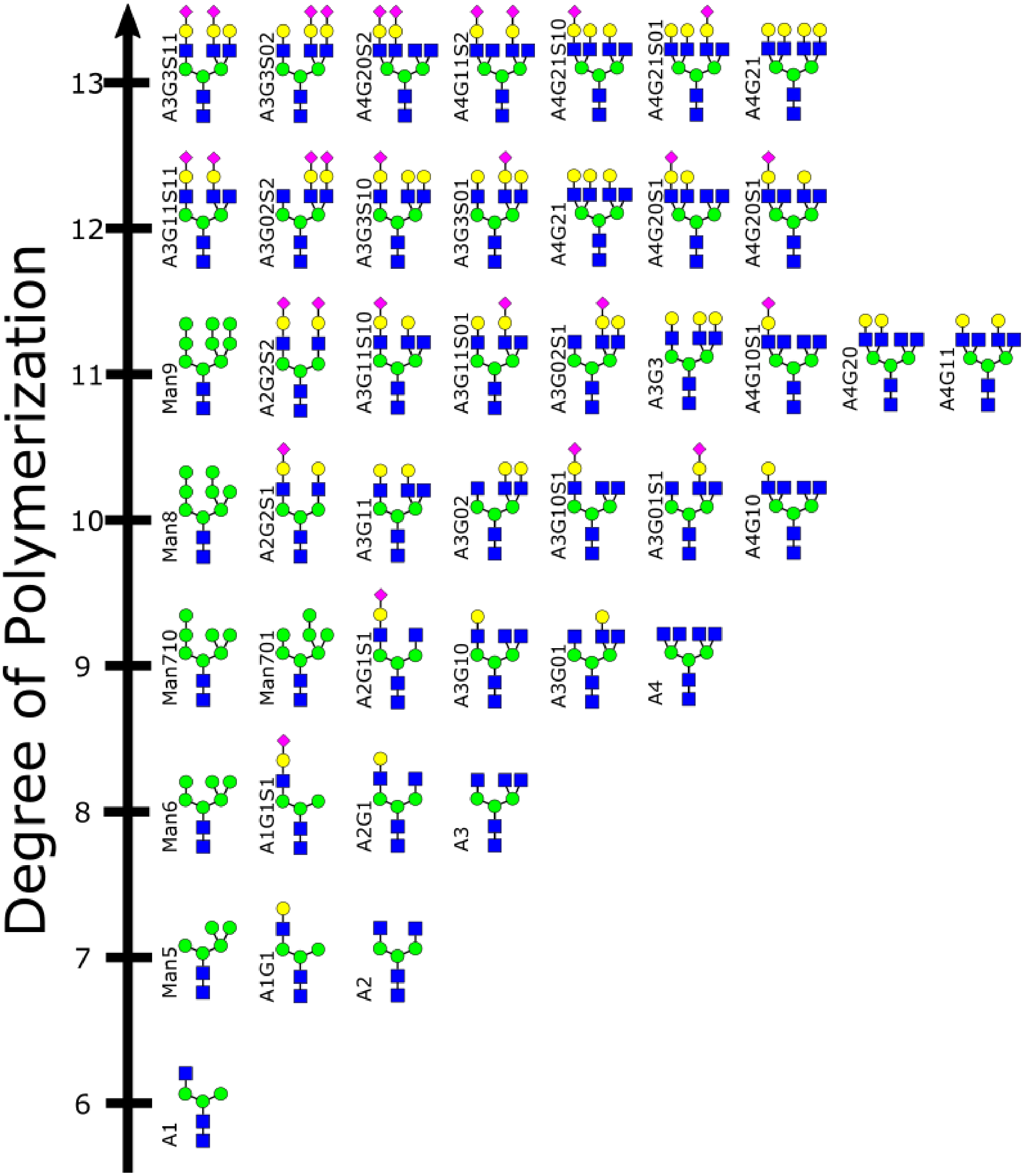


Figure S2: List of all the glycans considered in this study. The vertical axis gives the degree of polymerization, which is the number of sugar monomers connected to form the carbohydrate chain.

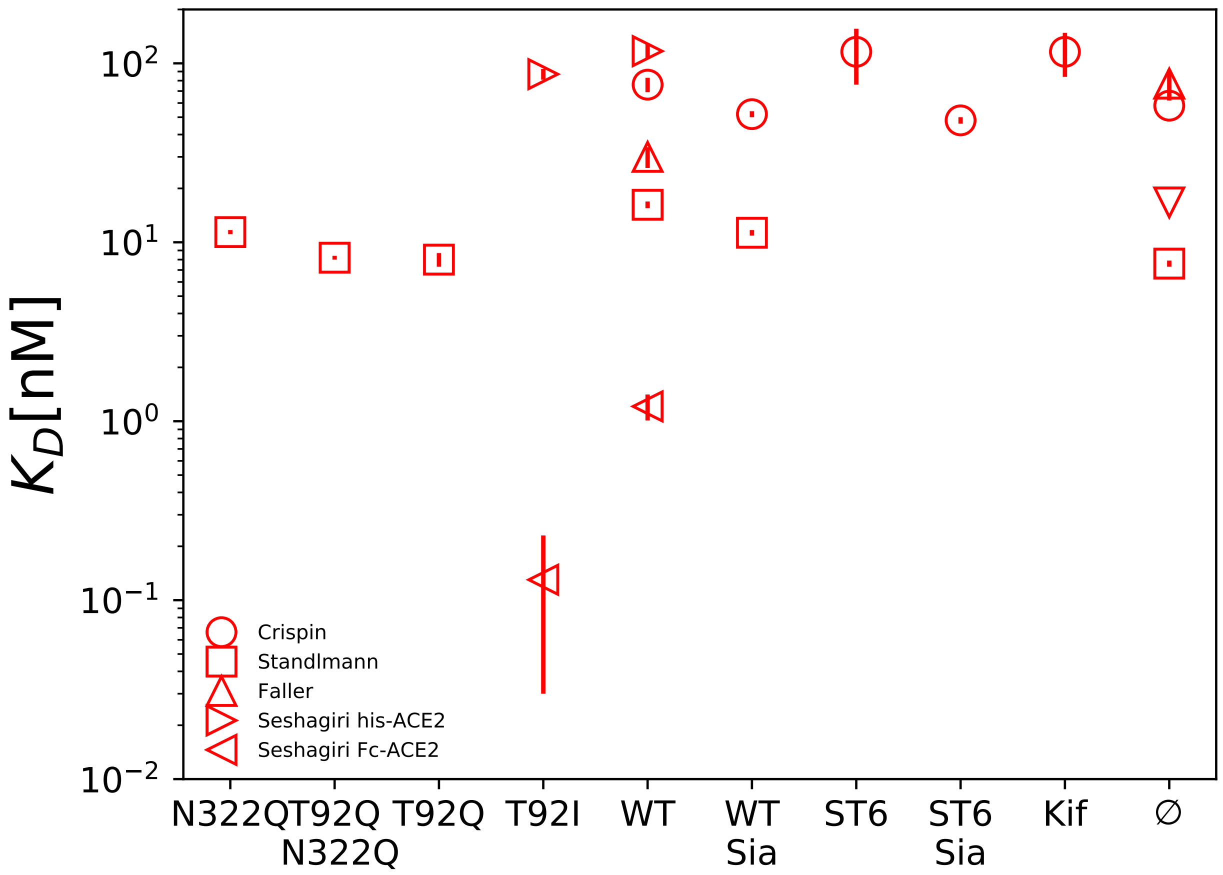


Figure S3: Experimental data analyzed in this study. The y-axis (in log-scale) refers to the dissociation constant of the various ACE2 glycoforms reported in the x-axis. For details, see caption to Fig. 3 in the main text.

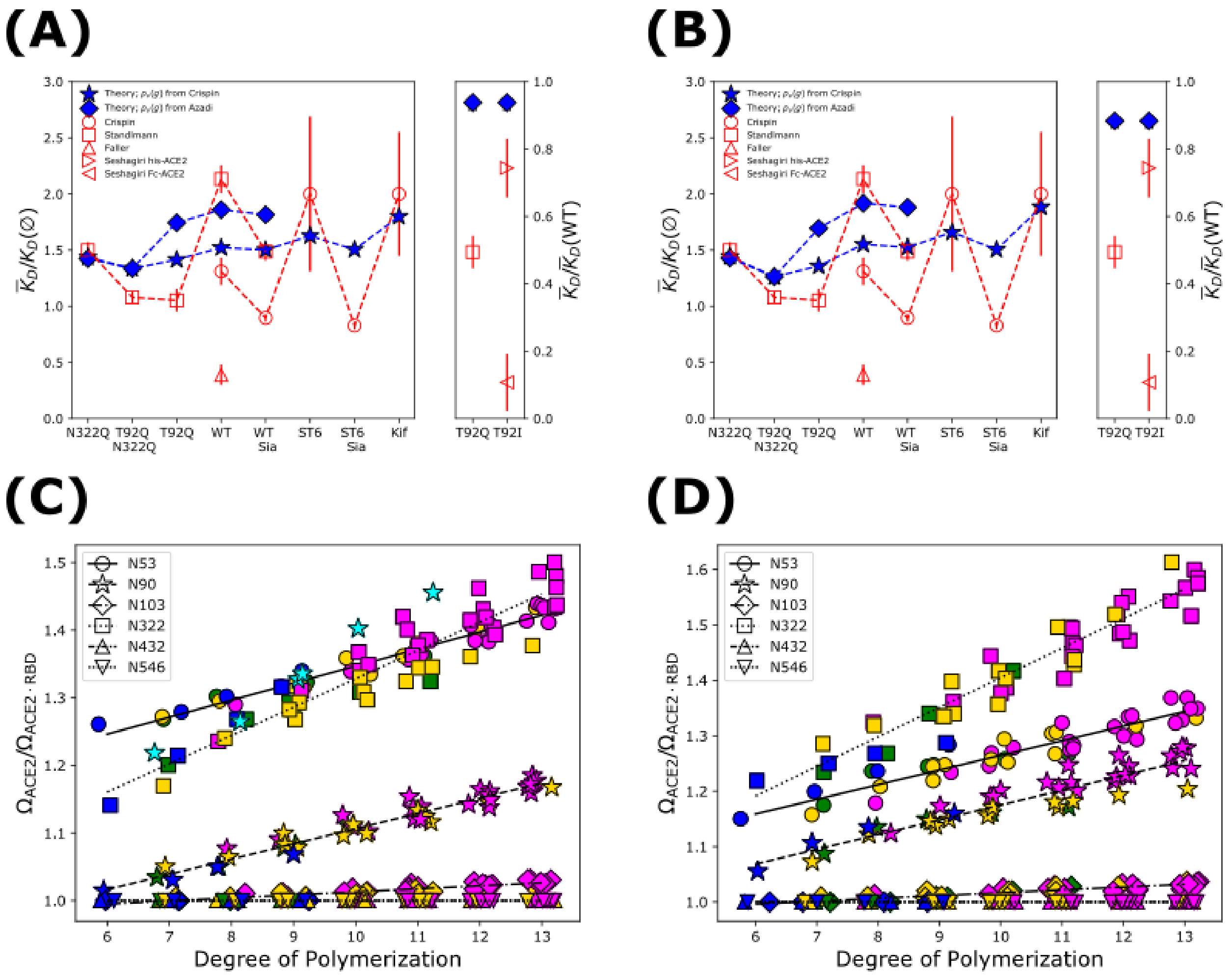


Figure S4: Robustness of the results over changes of the lattice spacing. The blue curves show the model dissociation constants for (A) $a = 5.0$ Å, (B) $a = 6.0$ Å. Panels (C) and (D) illustrate how the dependence of the dissociation constant on the length of the sugar chains changes when $a = 5.0$ Å (A) or $a = 6.0$ Å (B). In panel (C), the cyan stars indicate the contribution of the N90 oligosaccharide for the mutant ACE2 K26A (see Fig. S5). This calculation was performed using only the mannose-rich-type glycans (colored in green when referring to the WT ACE2).

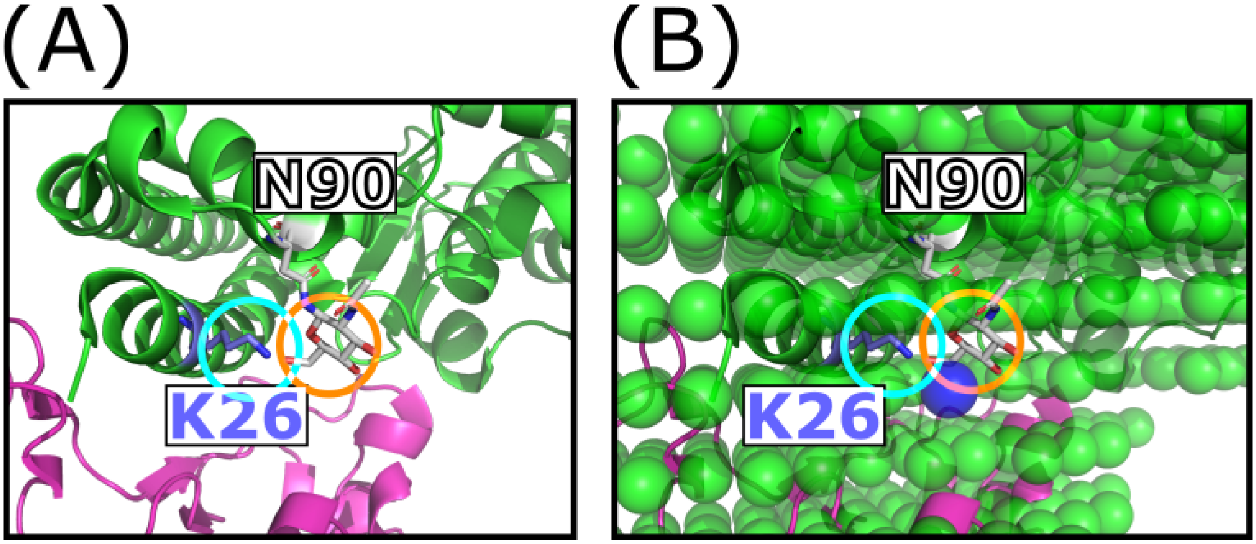


Figure S5: The role of ACE2 K26. (A) PDB ID: 6LZG is shown in cartoon (ACE2 in green, RBD in magenta). ACE2 residues K26 (blue) and N90 (white) are shown in stick mode, together with the first sugar GlcNAc of the N90 glycan (orange circle). The tip of the K26 side chain is highlighted with a cyan circle, and forms a wedge between the RBD and the N90 oligosaccharide. (B) Same as (A), but overlaid with the discretized representation of the protein complex, shown as green and blue spheres indicating the lattice sites occupied by the protein. The blue sphere is missing in the K26A mutant, which is obtained by removing the side chain of K26 after the beta carbon.

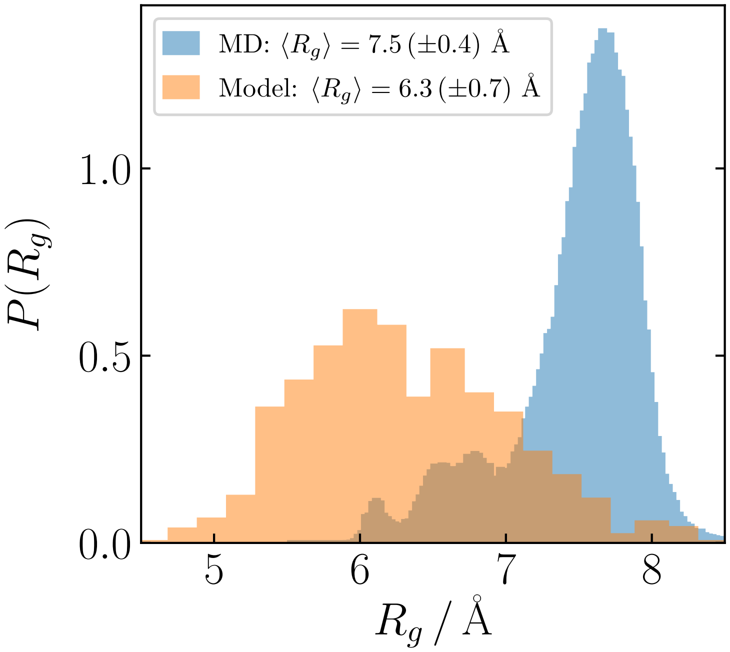


Figure S6: Comparison between the distributions of the radius of gyration computed from the MD simulation and the lattice model.

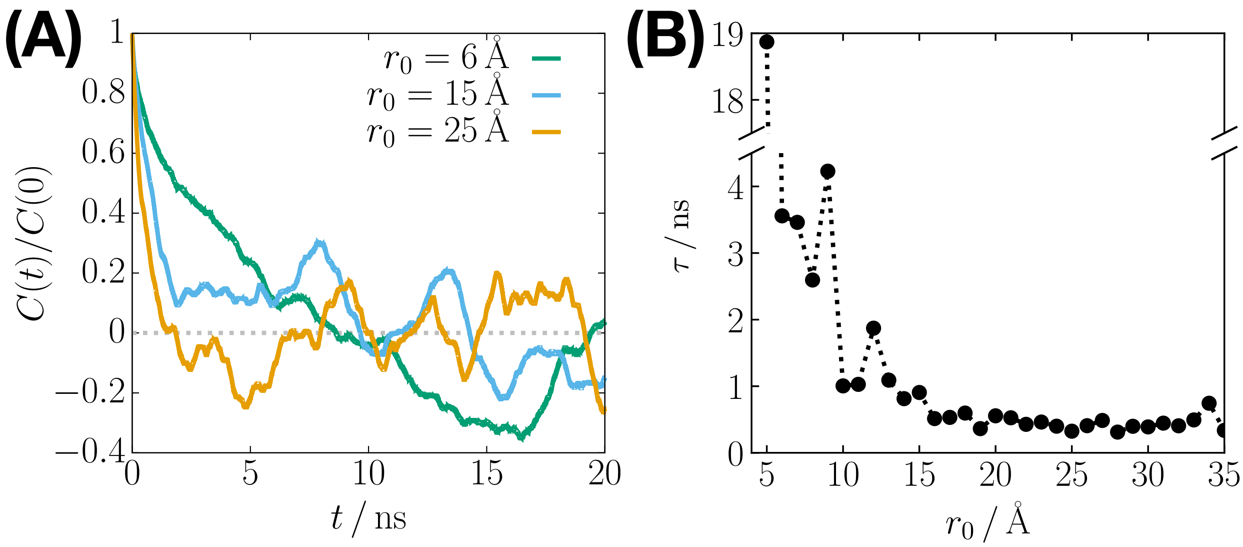


Figure S7: Validation of the observation time for the umbrella sampling. (A) Correlation function for the relative orientation between a pair of glycans [Eq. (S12)], at different equilibrium distance, $r_0$, of the biasing potential. (B) Correlation time as a function of $r_0$, where the breakage in y-axis is used to show the outlier for $r_0 = 5$ .

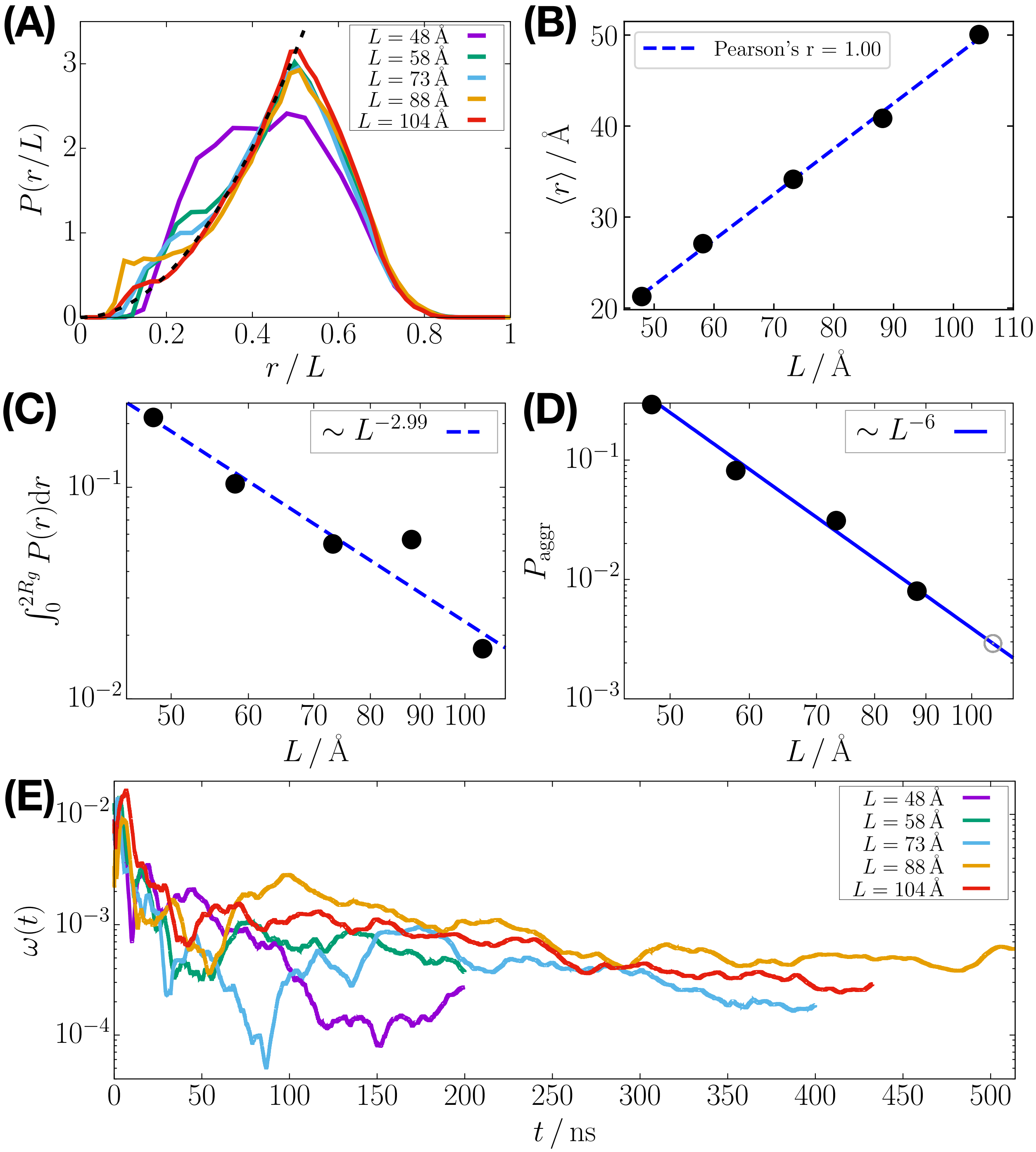


Figure S8: MD simulations reveal that glycans do not aggregate in water. (A) Probability distribution of the glycan pair distance, $r$, normalized by the length of the cubic simulation box, $L$. The black dashed line represents $4\pi(r/L)^2$. (B) Ensemble-averaged pair distance between the glycans as a function of $L$. The best linear fit is shown by the dashed line. (C) Log-log plot of the contact probability for a glycan pair as a function of $L$. The dashed line is the best linear fit. (D) Log-log plot of the aggregation probability [Eq. (S14)] as a function of $L$, where the blue solid line is the scaling relation expected for an ideal solution. The gray circle indicates the missing data point for $L = 10.4$ nm because $P_{\mathrm{aggr}} = 0$ for the given statistics. (E) Ergodic measures [Eq. (S13)] calculated using the trajectories for different simulation box sizes. In all cases, $\omega(t)$ is small at long times.

Table S1: Degree of polymerization and number of self-avoiding conformations for all of the glycans.

| glycan | degree of polymerization | number of conformations |
|---|---|---|
| Man5 | 7 | 8848 |
| Man6 | 8 | 40048 |
| Man710 | 9 | 187200 |
| Man701 | 9 | 186060 |
| Man8 | 10 | 855872 |
| Man9 | 11 | 3756976 |
| A1 | 6 | 2276 |
| A1G1 | 7 | 10532 |
| A1G1S1 | 8 | 50980 |
| A2 | 7 | 10684 |
| A2G1 | 8 | 49684 |
| A2G1S1 | 9 | 236592 |
| A2G2 | 9 | 227172 |
| A2G2S1 | 10 | 1085804 |
| A2G2S2 | 11 | 5144772 |
| A3 | 8 | 40048 |
| A3G10 | 9 | 187200 |
| A3G10S1 | 10 | 877952 |
| A3G01 | 9 | 186060 |
| A3G01S1 | 10 | 855116 |
| A3G11 | 10 | 855872 |
| A3G11S10 | 11 | 4028672 |
| A3G11S01 | 11 | 3947040 |
| A3G11S11 | 12 | 18422172 |
| A3G02 | 10 | 830840 |
| A3G02S1 | 11 | 3838360 |
| A3G02S2 | 12 | 17417496 |
| A3G3 | 11 | 3756976 |
| A3G3S10 | 12 | 17750272 |
| A3G3S01 | 12 | 17417496 |
| A3G3S11 | 13 | 81591860 |
| A3G3S02 | 13 | 79305352 |
| A4 | 9 | 144192 |
| A4G10 | 10 | 672912 |
| A4G10S1 | 11 | 3045552 |
| A4G20 | 11 | 3019200 |
| A4G20S1 | 12 | 13730648 |
| A4G20S2 | 13 | 61299296 |
| A4G11 | 11 | 3092408 |
| A4G11S10 | 12 | 14042984 |
| A4G11S11 | 13 | 63242600 |
| A4G21 | 12 | 13648912 |
| A4G21S10 | 13 | 62283736 |
| A4G21S01 | 13 | 62193320 |
| A4G22 | 13 | 59190432 |

Table S2: Values of $p_\nu(g)$ for the WT experiments of Crispin and coworkers.[S2]

| glycan | N53 | N90 | N103 | N322 |
|---|---|---|---|---|
| Empty | 0.0 | 0.0 | 0.03 | 0.72 |
| Man5 | 0.0 | 0.0 | 0.0 | 0.06 |
| Man6 | 0.0 | 0.04 | 0.0 | 0.0 |
| Man710 | 0.0 | 0.0 | 0.0 | 0.0 |
| Man701 | 0.0 | 0.0 | 0.0 | 0.0 |
| Man8 | 0.0 | 0.0 | 0.0 | 0.0 |
| Man9 | 0.0 | 0.0 | 0.0 | 0.0 |
| A1 | 0.0 | 0.01 | 0.24 | 0.11 |
| A1G1 | 0.02 | 0.01 | 0.02 | 0.0 |
| A1G1S1 | 0.0 | 0.0 | 0.0 | 0.0 |
| A2 | 0.127 | 0.27 | 0.1 | 0.0 |
| A2G1 | 0.044 | 0.15 | 0.065 | 0.005 |
| A2G2 | 0.044 | 0.15 | 0.065 | 0.005 |
| A2G1S1 | 0.026 | 0.017 | 0.107 | 0.0 |
| A2G2S1 | 0.026 | 0.017 | 0.107 | 0.0 |
| A2G2S2 | 0.026 | 0.017 | 0.107 | 0.0 |
| A3 | 0.284 | 0.08 | 0.05 | 0.06 |
| A3G10 | 0.031 | 0.02 | 0.022 | 0.002 |
| A3G01 | 0.031 | 0.02 | 0.022 | 0.002 |
| A3G11 | 0.031 | 0.02 | 0.022 | 0.002 |
| A3G02 | 0.031 | 0.02 | 0.022 | 0.002 |
| A3G3 | 0.031 | 0.02 | 0.022 | 0.002 |
| A3G10S1 | 0.0 | 0.011 | 0.0 | 0.0 |
| A3G01S1 | 0.0 | 0.011 | 0.0 | 0.0 |
| A3G11S10 | 0.0 | 0.011 | 0.0 | 0.0 |
| A3G11S01 | 0.0 | 0.011 | 0.0 | 0.0 |
| A3G11S11 | 0.0 | 0.011 | 0.0 | 0.0 |
| A3G02S1 | 0.0 | 0.011 | 0.0 | 0.0 |
| A3G02S2 | 0.0 | 0.011 | 0.0 | 0.0 |
| A3G3S10 | 0.0 | 0.011 | 0.0 | 0.0 |
| A3G3S01 | 0.0 | 0.011 | 0.0 | 0.0 |
| A3G3S11 | 0.0 | 0.011 | 0.0 | 0.0 |
| A3G3S02 | 0.0 | 0.011 | 0.0 | 0.0 |
| A4 | 0.098 | 0.0 | 0.0 | 0.0 |
| A4G10 | 0.016 | 0.003 | 0.0 | 0.005 |
| A4G10 | 0.016 | 0.003 | 0.0 | 0.005 |
| A4G11 | 0.016 | 0.003 | 0.0 | 0.005 |
| A4G20 | 0.016 | 0.003 | 0.0 | 0.005 |
| A4G21 | 0.016 | 0.003 | 0.0 | 0.005 |
| A4G22 | 0.016 | 0.003 | 0.0 | 0.005 |
| A4G10S1 | 0.007 | 0.0 | 0.0 | 0.0 |
| A4G20S1 | 0.007 | 0.0 | 0.0 | 0.0 |
| A4G20S2 | 0.007 | 0.0 | 0.0 | 0.0 |
| A4G11S10 | 0.007 | 0.0 | 0.0 | 0.0 |
| A4G11S11 | 0.007 | 0.0 | 0.0 | 0.0 |
| A4G21S10 | 0.007 | 0.0 | 0.0 | 0.0 |
| A4G21S01 | 0.007 | 0.0 | 0.0 | 0.0 |

Table S3: Values of $p_\nu(g)$ for the WT experiments of Crispin and coworkers[S2] plus sialidase.

| glycan | N53 | N90 | N103 | N322 |
|---|---|---|---|---|
| Empty | 0.0 | 0.0 | 0.03 | 0.72 |
| Man5 | 0.0 | 0.0 | 0.0 | 0.06 |
| Man6 | 0.0 | 0.04 | 0.0 | 0.0 |
| Man710 | 0.0 | 0.0 | 0.0 | 0.0 |
| Man701 | 0.0 | 0.0 | 0.0 | 0.0 |
| Man8 | 0.0 | 0.0 | 0.0 | 0.0 |
| Man9 | 0.0 | 0.0 | 0.0 | 0.0 |
| A1 | 0.0 | 0.01 | 0.24 | 0.11 |
| A1G1 | 0.02 | 0.01 | 0.02 | 0.0 |
| A1G1S1 | 0.0 | 0.0 | 0.0 | 0.0 |
| A2 | 0.127 | 0.27 | 0.1 | 0.0 |
| A2G1 | 0.083 | 0.175 | 0.225 | 0.005 |
| A2G2 | 0.083 | 0.175 | 0.225 | 0.005 |
| A2G1S1 | 0.0 | 0.0 | 0.0 | 0.0 |
| A2G2S1 | 0.0 | 0.0 | 0.0 | 0.0 |
| A2G2S2 | 0.0 | 0.0 | 0.0 | 0.0 |
| A3 | 0.284 | 0.08 | 0.05 | 0.06 |
| A3G10 | 0.031 | 0.044 | 0.022 | 0.002 |
| A3G01 | 0.031 | 0.044 | 0.022 | 0.002 |
| A3G11 | 0.031 | 0.044 | 0.022 | 0.002 |
| A3G02 | 0.031 | 0.044 | 0.022 | 0.002 |
| A3G3 | 0.031 | 0.044 | 0.022 | 0.002 |
| A3G10S1 | 0.0 | 0.0 | 0.0 | 0.0 |
| A3G01S1 | 0.0 | 0.0 | 0.0 | 0.0 |
| A3G11S10 | 0.0 | 0.0 | 0.0 | 0.0 |
| A3G11S01 | 0.0 | 0.0 | 0.0 | 0.0 |
| A3G11S11 | 0.0 | 0.0 | 0.0 | 0.0 |
| A3G02S1 | 0.0 | 0.0 | 0.0 | 0.0 |
| A3G02S2 | 0.0 | 0.0 | 0.0 | 0.0 |
| A3G3S10 | 0.0 | 0.0 | 0.0 | 0.0 |
| A3G3S01 | 0.0 | 0.0 | 0.0 | 0.0 |
| A3G3S11 | 0.0 | 0.0 | 0.0 | 0.0 |
| A3G3S02 | 0.0 | 0.0 | 0.0 | 0.0 |
| A4 | 0.098 | 0.0 | 0.0 | 0.0 |
| A4G10 | 0.025 | 0.003 | 0.0 | 0.005 |
| A4G10 | 0.025 | 0.003 | 0.0 | 0.005 |
| A4G11 | 0.025 | 0.003 | 0.0 | 0.005 |
| A4G20 | 0.025 | 0.003 | 0.0 | 0.005 |
| A4G21 | 0.025 | 0.003 | 0.0 | 0.005 |
| A4G22 | 0.025 | 0.003 | 0.0 | 0.005 |
| A4G10S1 | 0.0 | 0.0 | 0.0 | 0.0 |
| A4G20S1 | 0.0 | 0.0 | 0.0 | 0.0 |
| A4G20S2 | 0.0 | 0.0 | 0.0 | 0.0 |
| A4G11S10 | 0.0 | 0.0 | 0.0 | 0.0 |
| A4G11S11 | 0.0 | 0.0 | 0.0 | 0.0 |
| A4G21S10 | 0.0 | 0.0 | 0.0 | 0.0 |
| A4G21S01 | 0.0 | 0.0 | 0.0 | 0.0 |

Table S4: Values of $p_\nu(g)$ for the +ST6 experiments of Crispin and coworkers.[S2]

| glycan | N53 | N90 | N103 | N322 |
|---|---|---|---|---|
| Empty | 0.0 | 0.0 | 0.0 | 0.86 |
| Man5 | 0.0 | 0.0 | 0.0 | 0.0 |
| Man6 | 0.0 | 0.0 | 0.0 | 0.0 |
| Man710 | 0.0 | 0.0 | 0.0 | 0.0 |
| Man701 | 0.0 | 0.0 | 0.0 | 0.0 |
| Man8 | 0.0 | 0.0 | 0.0 | 0.0 |
| Man9 | 0.0 | 0.0 | 0.0 | 0.0 |
| A1 | 0.011 | 0.01 | 0.152 | 0.0 |
| A1G1 | 0.0 | 0.0 | 0.0 | 0.0 |
| A1G1S1 | 0.0 | 0.01 | 0.0 | 0.0 |
| A2 | 0.138 | 0.1 | 0.071 | 0.03 |
| A2G1 | 0.0 | 0.005 | 0.02 | 0.0 |
| A2G2 | 0.0 | 0.005 | 0.02 | 0.0 |
| A2G1S1 | 0.0 | 0.243 | 0.047 | 0.0 |
| A2G2S1 | 0.0 | 0.243 | 0.047 | 0.0 |
| A2G2S2 | 0.0 | 0.243 | 0.047 | 0.0 |
| A3 | 0.011 | 0.01 | 0.0 | 0.0 |
| A3G10 | 0.002 | 0.0 | 0.012 | 0.0 |
| A3G01 | 0.002 | 0.0 | 0.012 | 0.0 |
| A3G11 | 0.002 | 0.0 | 0.012 | 0.0 |
| A3G02 | 0.002 | 0.0 | 0.012 | 0.0 |
| A3G3 | 0.002 | 0.0 | 0.012 | 0.0 |
| A3G10S1 | 0.071 | 0.012 | 0.048 | 0.001 |
| A3G01S1 | 0.071 | 0.012 | 0.048 | 0.001 |
| A3G11S10 | 0.071 | 0.012 | 0.048 | 0.001 |
| A3G11S01 | 0.071 | 0.012 | 0.048 | 0.001 |
| A3G11S11 | 0.071 | 0.012 | 0.048 | 0.001 |
| A3G02S1 | 0.071 | 0.012 | 0.048 | 0.001 |
| A3G02S2 | 0.071 | 0.012 | 0.048 | 0.001 |
| A3G3S10 | 0.071 | 0.012 | 0.048 | 0.001 |
| A3G3S01 | 0.071 | 0.012 | 0.048 | 0.001 |
| A3G3S11 | 0.071 | 0.012 | 0.048 | 0.001 |
| A3G3S02 | 0.071 | 0.012 | 0.048 | 0.001 |
| A4 | 0.0 | 0.0 | 0.0 | 0.0 |
| A4G10 | 0.0 | 0.0 | 0.002 | 0.0 |
| A4G10 | 0.0 | 0.0 | 0.002 | 0.0 |
| A4G11 | 0.0 | 0.0 | 0.002 | 0.0 |
| A4G20 | 0.0 | 0.0 | 0.002 | 0.0 |
| A4G21 | 0.0 | 0.0 | 0.002 | 0.0 |
| A4G22 | 0.0 | 0.0 | 0.002 | 0.0 |
| A4G10S1 | 0.008 | 0.0 | 0.0 | 0.014 |
| A4G20S1 | 0.008 | 0.0 | 0.0 | 0.014 |
| A4G20S2 | 0.008 | 0.0 | 0.0 | 0.014 |
| A4G11S10 | 0.008 | 0.0 | 0.0 | 0.014 |
| A4G11S11 | 0.008 | 0.0 | 0.0 | 0.014 |
| A4G21S10 | 0.008 | 0.0 | 0.0 | 0.014 |
| A4G21S01 | 0.008 | 0.0 | 0.0 | 0.014 |

Table S5: Values of $p_\nu(g)$ for the +ST6 experiments of Crispin and coworkers[S2] plus sialidase.

| glycan | N53 | N90 | N103 | N322 |
|---|---|---|---|---|
| Empty | 0.0 | 0.0 | 0.0 | 0.86 |
| Man5 | 0.0 | 0.0 | 0.0 | 0.0 |
| Man6 | 0.0 | 0.0 | 0.0 | 0.0 |
| Man710 | 0.0 | 0.0 | 0.0 | 0.0 |
| Man701 | 0.0 | 0.0 | 0.0 | 0.0 |
| Man8 | 0.0 | 0.0 | 0.0 | 0.0 |
| Man9 | 0.0 | 0.0 | 0.0 | 0.0 |
| A1 | 0.01 | 0.01 | 0.152 | 0.0 |
| A1G1 | 0.0 | 0.01 | 0.0 | 0.0 |
| A1G1S1 | 0.0 | 0.0 | 0.0 | 0.0 |
| A2 | 0.168 | 0.1 | 0.071 | 0.03 |
| A2G1 | 0.015 | 0.37 | 0.091 | 0.0 |
| A2G2 | 0.015 | 0.37 | 0.091 | 0.0 |
| A2G1S1 | 0.0 | 0.0 | 0.0 | 0.0 |
| A2G2S1 | 0.0 | 0.0 | 0.0 | 0.0 |
| A2G2S2 | 0.0 | 0.0 | 0.0 | 0.0 |
| A3 | 0.01 | 0.01 | 0.0 | 0.0 |
| A3G10 | 0.147 | 0.026 | 0.117 | 0.002 |
| A3G01 | 0.147 | 0.026 | 0.117 | 0.002 |
| A3G11 | 0.147 | 0.026 | 0.117 | 0.002 |
| A3G02 | 0.147 | 0.026 | 0.117 | 0.002 |
| A3G3 | 0.147 | 0.026 | 0.117 | 0.002 |
| A3G10S1 | 0.0 | 0.0 | 0.0 | 0.0 |
| A3G01S1 | 0.0 | 0.0 | 0.0 | 0.0 |
| A3G11S10 | 0.0 | 0.0 | 0.0 | 0.0 |
| A3G11S01 | 0.0 | 0.0 | 0.0 | 0.0 |
| A3G11S11 | 0.0 | 0.0 | 0.0 | 0.0 |
| A3G02S1 | 0.0 | 0.0 | 0.0 | 0.0 |
| A3G02S2 | 0.0 | 0.0 | 0.0 | 0.0 |
| A3G3S10 | 0.0 | 0.0 | 0.0 | 0.0 |
| A3G3S01 | 0.0 | 0.0 | 0.0 | 0.0 |
| A3G3S11 | 0.0 | 0.0 | 0.0 | 0.0 |
| A3G3S02 | 0.0 | 0.0 | 0.0 | 0.0 |
| A4 | 0.0 | 0.0 | 0.0 | 0.0 |
| A4G10 | 0.008 | 0.0 | 0.002 | 0.017 |
| A4G10 | 0.008 | 0.0 | 0.002 | 0.017 |
| A4G11 | 0.008 | 0.0 | 0.002 | 0.017 |
| A4G20 | 0.008 | 0.0 | 0.002 | 0.017 |
| A4G21 | 0.008 | 0.0 | 0.002 | 0.017 |
| A4G22 | 0.008 | 0.0 | 0.002 | 0.017 |
| A4G10S1 | 0.0 | 0.0 | 0.0 | 0.0 |
| A4G20S1 | 0.0 | 0.0 | 0.0 | 0.0 |
| A4G20S2 | 0.0 | 0.0 | 0.0 | 0.0 |
| A4G11S10 | 0.0 | 0.0 | 0.0 | 0.0 |
| A4G11S11 | 0.0 | 0.0 | 0.0 | 0.0 |
| A4G21S10 | 0.0 | 0.0 | 0.0 | 0.0 |
| A4G21S01 | 0.0 | 0.0 | 0.0 | 0.0 |

Table S6: Values of $p_\nu(g)$ for the +Kif experiments of Crispin and coworkers.[S2]

| glycan | N53 | N90 | N103 | N322 |
|---|---|---|---|---|
| Empty | 0.0 | 0.0 | 0.0 | 0.51 |
| Man5 | 0.0 | 0.0 | 0.0 | 0.0 |
| Man6 | 0.0 | 0.0 | 0.0 | 0.0 |
| Man710 | 0.0 | 0.09 | 0.1 | 0.0 |
| Man701 | 0.0 | 0.09 | 0.1 | 0.0 |
| Man8 | 0.0 | 0.01 | 0.0 | 0.0 |
| Man9 | 1.0 | 0.81 | 0.79 | 0.49 |
| A1 | 0.0 | 0.0 | 0.0 | 0.0 |
| A1G1 | 0.0 | 0.0 | 0.0 | 0.0 |
| A1G1S1 | 0.0 | 0.0 | 0.0 | 0.0 |
| A2 | 0.0 | 0.0 | 0.01 | 0.0 |
| A2G1 | 0.0 | 0.0 | 0.0 | 0.0 |
| A2G2 | 0.0 | 0.0 | 0.0 | 0.0 |
| A2G1S1 | 0.0 | 0.0 | 0.0 | 0.0 |
| A2G2S1 | 0.0 | 0.0 | 0.0 | 0.0 |
| A2G2S2 | 0.0 | 0.0 | 0.0 | 0.0 |
| A3 | 0.0 | 0.0 | 0.0 | 0.0 |
| A3G10 | 0.0 | 0.0 | 0.0 | 0.0 |
| A3G01 | 0.0 | 0.0 | 0.0 | 0.0 |
| A3G11 | 0.0 | 0.0 | 0.0 | 0.0 |
| A3G02 | 0.0 | 0.0 | 0.0 | 0.0 |
| A3G3 | 0.0 | 0.0 | 0.0 | 0.0 |
| A3G10S1 | 0.0 | 0.0 | 0.0 | 0.0 |
| A3G01S1 | 0.0 | 0.0 | 0.0 | 0.0 |
| A3G11S10 | 0.0 | 0.0 | 0.0 | 0.0 |
| A3G11S01 | 0.0 | 0.0 | 0.0 | 0.0 |
| A3G11S11 | 0.0 | 0.0 | 0.0 | 0.0 |
| A3G02S1 | 0.0 | 0.0 | 0.0 | 0.0 |
| A3G02S2 | 0.0 | 0.0 | 0.0 | 0.0 |
| A3G3S10 | 0.0 | 0.0 | 0.0 | 0.0 |
| A3G3S01 | 0.0 | 0.0 | 0.0 | 0.0 |
| A3G3S11 | 0.0 | 0.0 | 0.0 | 0.0 |
| A3G3S02 | 0.0 | 0.0 | 0.0 | 0.0 |
| A4 | 0.0 | 0.0 | 0.0 | 0.0 |
| A4G10 | 0.0 | 0.0 | 0.0 | 0.0 |
| A4G10 | 0.0 | 0.0 | 0.0 | 0.0 |
| A4G11 | 0.0 | 0.0 | 0.0 | 0.0 |
| A4G20 | 0.0 | 0.0 | 0.0 | 0.0 |
| A4G21 | 0.0 | 0.0 | 0.0 | 0.0 |
| A4G22 | 0.0 | 0.0 | 0.0 | 0.0 |
| A4G10S1 | 0.0 | 0.0 | 0.0 | 0.0 |
| A4G20S1 | 0.0 | 0.0 | 0.0 | 0.0 |
| A4G20S2 | 0.0 | 0.0 | 0.0 | 0.0 |
| A4G11S10 | 0.0 | 0.0 | 0.0 | 0.0 |
| A4G11S11 | 0.0 | 0.0 | 0.0 | 0.0 |
| A4G21S10 | 0.0 | 0.0 | 0.0 | 0.0 |
| A4G21S01 | 0.0 | 0.0 | 0.0 | 0.0 |

Table S7: Values of $p_\nu(g)$ for the WT experiments of Azadi and coworkers.[S4]

| glycan | N53 | N90 | N103 | N322 |
|---|---|---|---|---|
| A1 | 0.0 | 0.044 | 0.018 | 0.0 |
| A2 | 0.122 | 0.23 | 0.578 | 0.099 |
| A2G1 | 0.03 | 0.43 | 0.302 | 0.0 |
| A2G1S1 | 0.234 | 0.167 | 0.0 | 0.0 |
| A2G2 | 0.0 | 0.018 | 0.073 | 0.0 |
| A3 | 0.067 | 0.0 | 0.0 | 0.417 |
| A3G10 | 0.027 | 0.0 | 0.0 | 0.072 |
| A3G01 | 0.027 | 0.0 | 0.0 | 0.072 |
| A3G11 | 0.0 | 0.0 | 0.0 | 0.0 |
| A3G02 | 0.0 | 0.0 | 0.0 | 0.0 |
| A3G3 | 0.0 | 0.0 | 0.0 | 0.0 |
| A4 | 0.042 | 0.0 | 0.0 | 0.013 |
| A4G10 | 0.048 | 0.0 | 0.0 | 0.011 |
| A2G2S1 | 0.055 | 0.11 | 0.029 | 0.0 |
| A3G11 | 0.0 | 0.0 | 0.0 | 0.042 |
| A3G10S1 | 0.096 | 0.0 | 0.0 | 0.033 |
| A3G01S1 | 0.096 | 0.0 | 0.0 | 0.033 |
| A3G11S10 | 0.01 | 0.0 | 0.0 | 0.027 |
| A3G11S01 | 0.01 | 0.0 | 0.0 | 0.027 |
| A3G02S1 | 0.01 | 0.0 | 0.0 | 0.027 |
| A3G3S10 | 0.0 | 0.0 | 0.0 | 0.021 |
| A3G3S01 | 0.0 | 0.0 | 0.0 | 0.021 |
| A3G3S11 | 0.0 | 0.0 | 0.0 | 0.009 |
| A3G3S02 | 0.0 | 0.0 | 0.0 | 0.009 |
| A4G20 | 0.0 | 0.0 | 0.0 | 0.0 |
| A4G11 | 0.0 | 0.0 | 0.0 | 0.0 |
| A4G21 | 0.0 | 0.0 | 0.0 | 0.0 |
| A4G21 | 0.0 | 0.0 | 0.0 | 0.0 |
| A4G4 | 0.0 | 0.0 | 0.0 | 0.0 |
| A4G10S1 | 0.073 | 0.0 | 0.0 | 0.0 |
| A4G20S1 | 0.018 | 0.0 | 0.0 | 0.005 |
| A4G11S10 | 0.018 | 0.0 | 0.0 | 0.005 |
| A4G20S2 | 0.0 | 0.0 | 0.0 | 0.005 |
| A4G11S11 | 0.0 | 0.0 | 0.0 | 0.005 |
| A4G21S10 | 0.009 | 0.0 | 0.0 | 0.005 |
| A4G21S01 | 0.009 | 0.0 | 0.0 | 0.005 |
| A4G20S2 | 0.0 | 0.0 | 0.0 | 0.006 |
| A4G11S11 | 0.0 | 0.0 | 0.0 | 0.006 |
| A4G20S2 | 0.0 | 0.0 | 0.0 | 0.009 |
| A4G11S11 | 0.0 | 0.0 | 0.0 | 0.009 |

Table S8: Values of $p_\nu(g)$ for the WT experiments of Azadi and coworkers[S4] plus sialidase.

| glycan | N53 | N90 | N103 | N322 |
|---|---|---|---|---|
| A1 | 0.0 | 0.044 | 0.018 | 0.0 |
| A2 | 0.122 | 0.23 | 0.578 | 0.097 |
| A2G1 | 0.03 | 0.598 | 0.302 | 0.0 |
| A2G1S1 | 0.234 | 0.0 | 0.0 | 0.0 |
| A2G2 | 0.0 | 0.128 | 0.101 | 0.0 |
| A3 | 0.067 | 0.0 | 0.0 | 0.409 |
| A3G10 | 0.027 | 0.0 | 0.0 | 0.104 |
| A3G01 | 0.027 | 0.0 | 0.0 | 0.104 |
| A3G11 | 0.0 | 0.0 | 0.0 | 0.04 |
| A3G02 | 0.0 | 0.0 | 0.0 | 0.04 |
| A3G3 | 0.0 | 0.0 | 0.0 | 0.079 |
| A4 | 0.042 | 0.0 | 0.0 | 0.013 |
| A4G10 | 0.048 | 0.0 | 0.0 | 0.011 |
| A2G2S1 | 0.055 | 0.0 | 0.0 | 0.0 |
| A3G11 | 0.0 | 0.0 | 0.0 | 0.041 |
| A3G10S1 | 0.096 | 0.0 | 0.0 | 0.0 |
| A3G01S1 | 0.096 | 0.0 | 0.0 | 0.0 |
| A3G11S10 | 0.01 | 0.0 | 0.0 | 0.0 |
| A3G11S01 | 0.01 | 0.0 | 0.0 | 0.0 |
| A3G02S1 | 0.01 | 0.0 | 0.0 | 0.0 |
| A3G3S10 | 0.0 | 0.0 | 0.0 | 0.0 |
| A3G3S01 | 0.0 | 0.0 | 0.0 | 0.0 |
| A3G3S11 | 0.0 | 0.0 | 0.0 | 0.0 |
| A3G3S02 | 0.0 | 0.0 | 0.0 | 0.0 |
| A4G20 | 0.0 | 0.0 | 0.0 | 0.011 |
| A4G11 | 0.0 | 0.0 | 0.0 | 0.011 |
| A4G21 | 0.0 | 0.0 | 0.0 | 0.0 |
| A4G21 | 0.0 | 0.0 | 0.0 | 0.022 |
| A4G4 | 0.0 | 0.0 | 0.0 | 0.019 |
| A4G10S1 | 0.073 | 0.0 | 0.0 | 0.0 |
| A4G20S1 | 0.018 | 0.0 | 0.0 | 0.0 |
| A4G11S10 | 0.018 | 0.0 | 0.0 | 0.0 |
| A4G20S2 | 0.0 | 0.0 | 0.0 | 0.0 |
| A4G11S11 | 0.0 | 0.0 | 0.0 | 0.0 |
| A4G21S10 | 0.009 | 0.0 | 0.0 | 0.0 |
| A4G21S01 | 0.009 | 0.0 | 0.0 | 0.0 |
| A4G20S2 | 0.0 | 0.0 | 0.0 | 0.0 |
| A4G11S11 | 0.0 | 0.0 | 0.0 | 0.0 |
| A4G20S2 | 0.0 | 0.0 | 0.0 | 0.0 |
| A4G11S11 | 0.0 | 0.0 | 0.0 | 0.0 |

(S5) de Gennes, P.-G. *Scaling Concepts in Polymer Physics*; Cornell University Press: Ithaca, NY, 1979.

(S6) Chen, C. L.; Hsu, J. C.; Lin, C. W.; Wang, C. H.; Tsai, M. H.; Wu, C. Y.; Wong, C. H.; Ma, C. *ACS Chemical Biology* **2017**, *12*, 1335–1345.

(S7) MacKerell, A. D.; Raman, E. P.; Guvench, O. *Journal of Physical Chemistry B* **2010**, *114*, 12981–12994.

(S8) Jorgensen, W. L.; Chandrasekhar, J.; Madura, J. D.; Impey, R. W.; Klein, M. L. *The Journal of Chemical Physics* **1983**, *79*, 926–935.

(S9) Humphrey, W.; Dalke, A.; Schulten, K. *Journal of Molecular Graphics* **1996**, *14*, 33–38.

(S10) Hoover, W. G. *Physical Review A* **1985**, *31*, 1695–1697.

(S11) Parrinello, M.; Rahman, A. *Journal of Applied Physics* **1981**, *52*, 7182–7190.

(S12) Darden, T.; York, D.; Pedersen, L. *The Journal of Chemical Physics* **1993**, *98*, 10089–10092.

(S13) Plimpton, S. *Journal of Computational Physics* **1995**, *117*, 1–19.

(S14) Kumar, S.; Bouzida, D.; Swendsen, R. H.; Kollman, P. A.; Rosenberg, J. M. *Journal of Computational Chemistry* **1992**, *13*, 1011–1021.

(S15) Thirumalai, D.; Mountain, R. D.; Kirkpatrick, T. R. *Physical Review A* **1989**, *39*, 3563–3574.

(S16) Thirumalai, D.; Mountain, R. D. *Physical Review A* **1990**, *42*, 4574–4587.

(S17) Straub, J. E.; Rashkin, A. B.; Thirumalai, D. *Journal of the American Chemical Society* **1994**, *116*, 2049–2063.
